# Survey of white-footed mice in Connecticut, USA reveals low SARS-CoV-2 seroprevalence and infection with divergent betacoronaviruses

**DOI:** 10.1101/2023.09.22.559030

**Authors:** Rebecca Earnest, Anne M. Hahn, Nicole M. Feriancek, Matthew Brandt, Renata B. Filler, Zhe Zhao, Mallery I. Breban, Chantal B.F. Vogels, Nicholas F.G. Chen, Robert T. Koch, Abbey J. Porzucek, Afeez Sodeinde, Alexa Garbiel, Claire Keanna, Hannah Litwak, Heidi R. Stuber, Jamie L. Cantoni, Virginia E. Pitzer, Ximena A. Olarte Castillo, Laura B. Goodman, Craig B. Wilen, Megan A. Linske, Scott C. Williams, Nathan D. Grubaugh

**Affiliations:** Department of Epidemiology of Microbial Diseases, Yale School of Public Health, New Haven, CT 06510, USA; Department of Laboratory Medicine, Yale School of Medicine, New Haven, CT 06520, USA; Department of Immunobiology, Yale School of Medicine, New Haven, CT 06520, USA; Department of Entomology, The Connecticut Agricultural Experiment Station, New Haven, CT 06511, USA; Department of Microbiology and Immunology, Cornell University College of Veterinary Medicine, Ithaca, NY 14853; Department of Public & Ecosystem Health, Cornell University College of Veterinary Medicine, Ithaca, NY 14853; Department of Environmental Science and Forestry, The Connecticut Agricultural Experiment Station, New Haven, CT 06511, USA; Department of Ecology and Evolutionary Biology, Yale University, New Haven, CT 06510, USA

**Author notes:** Co-first authors (Rebecca Earnest, Anne M. Hahn). Co-senior authors (Scott C. Williams, Nathan D. Grubaugh).

## Abstract

Diverse mammalian species display susceptibility to and infection with SARS-CoV-2. Potential SARS-CoV-2 spillback into rodents is understudied despite their host role for numerous zoonoses and human proximity. We assessed exposure and infection among white-footed mice (*Peromyscus leucopus*) in Connecticut, USA. We observed 1% (6/540) wild-type neutralizing antibody seroprevalence among 2020-2022 residential mice with no cross-neutralization of variants. We detected no SARS-CoV-2 infections via RT-qPCR, but identified non-SARS-CoV-2 betacoronavirus infections via pan-coronavirus PCR among 1% (5/468) of residential mice. Sequencing revealed two divergent betacoronaviruses, preliminarily named *Peromyscus coronavirus-1* and *-2*. Both belong to the *Betacoronavirus 1* species and are ∼90% identical to the closest known relative, *Porcine hemagglutinating encephalomyelitis virus*. Low SARS-CoV-2 seroprevalence suggests white-footed mice may not be sufficiently susceptible or exposed to SARS-CoV-2 to present a long-term human health risk. However, the discovery of divergent, non-SARS-CoV-2 betacoronaviruses expands the diversity of known rodent coronaviruses and further investigation is required to understand their transmission extent.

## Introduction

The COVID-19 pandemic likely began with an initial zoonotic spillover from an unknown animal species into humans, triggering unprecedented waves of infection^1^. In the intervening years, COVID-19 research necessarily focused on human-to-human transmission. However, there is another important dynamic to consider: SARS-CoV-2 transmission at the animal-human interface. Specifically, the risk posed by viral spillback (human-to-animal), sustained transmission in naive animal populations (animal-to-animal), and secondary spillover (spillback followed by animal-to-human) is currently unclear^2^. Outbreaks at the animal-human interface hold two important implications for long-term control. First, with the pandemic now in an endemic stage marked by less human-to-human transmission, animals could become an increasingly important source of new outbreaks. Second, transmission within new non-human host species could result in unexpected viral evolution, potentially yielding variants with properties such as increased virulence or enhanced immune escape^3^. Coupled with the challenges of viral surveillance in animal populations, concerning variants might not be detected until their emergence among humans.

For viral spillback to occur, a given animal species must be both susceptible and exposed to SARS-CoV-2. All vertebrates are potentially susceptible due to the highly conserved angiotensin-converting enzyme 2 (ACE2) receptor used by SARS-CoV-2 to enter host cells, although the degree of susceptibility varies based on how well the species’ ACE2 receptor binds to the virus spike protein receptor-binding domain (RBD)^4,5^. In addition, continually high levels of global human-to-human transmission have provided innumerable exposure opportunities over the past several years. As of August 2023, one database reported SARS-CoV-2 infections or exposures in 34 animal species across 39 countries^6^. Affected species include companion, zoo, farmed, and wild animals. If viral spillback occurred, only a subset of species may be capable of sufficiently high viral replication and infectious virus shedding, combined with suitable behaviors, to enable within-species transmission. In addition, sustained transmission within some species may depend on repeated viral spillback from humans^2^. The final dynamic to consider is that of potential secondary spillover. Few examples of secondary spillover of any pathogen have been reported, likely in part due to the many behavioral and host barriers a pathogen must overcome combined with the challenges of observing such dynamics^2^. Despite this, potentially due to broad susceptibility and substantial exposure opportunities, several instances of SARS-CoV-2 secondary spillover have been reported including from farmed mink (*Neovison vison*) to workers in the Netherlands, imported golden hamsters (*Mesocricetus auratus*) to pet shop workers in Hong Kong, and white-tailed deer (*Odocoileus virginianus*) to human contacts in Canada and the United States^7–10^. Not all animals are equally likely to experience viral spillback, sustained transmission, and/or secondary spillover. In this study, we focused on rodents due to their prominence as zoonotic disease reservoirs and proximity to humans^11^, posing the following question: is there evidence of SARS-CoV-2 spillback from humans or other animals into rodents in Connecticut (CT) and, if so, do we observe evidence of sustained transmission and secondary spillback?

To investigate, we selected the white-footed mouse (*Peromyscus leucopus*), a peridomestic species abundant throughout North America^12^. Experimental challenge studies show that two closely related species, the deer mouse (*Peromyscus maniculatus*) and California mouse (*Peromyscus californicus*), are capable of wild-type SARS-CoV-2 infection and direct transmission^13–16^. Among challenged *Peromyscine* species, only the California mouse displays clinical disease. Two studies have performed limited surveys of wild *Peromyscine* mice for SARS-CoV-2^17,18^. One study reported 17% (1/6) and 29% (4/14) neutralizing antibody seroprevalence among white-footed mice and deer mice, respectively^17^. The other found 4% rRT-PCR SARS-CoV-2 positivity for one of two gene targets among *Peromyscine* mice (2/50)^18^. The majority of other SARS-CoV-2 wild rodent studies focused on sewer rats, especially the Norway rat (*Rattus norvegicus*)^19–21^. A New York City study found that 16.5% of Norway rats were IgG or IgM positive and 5% RT-qPCR positivity among lung tissue samples^21^. SARS-CoV-2 surveillance in rodents remains highly limited despite these occasional detections. While rodents are the focus of our study, we also evaluated white-tailed deer, a species with extensive evidence of infection and exposure across the United States and Canada to provide a comparison^8,10,22–32^. Experimental challenge studies further confirm the SARS-CoV-2 infection and transmission capacity of deer, and note that the disease presents subclinically^33–35^.

We collected sera and swab samples from each species. We observed a 1% (6/540), 0% (0/69), and 7% (4/55) seroprevalence of wild-type SARS-CoV-2 neutralizing antibodies among residential white-footed mice, forested white-footed mice, and all deer, respectively. Seropositive deer most effectively neutralized wild-type SARS-CoV-2, with reduced or absent cross-neutralization of subsequent viral variants. While we did not detect active SARS-CoV-2 infections, we observed 1% (5/468), 0% (0/146), and 0% (0/31) pan-coronavirus PCR positivity among 2022 residential white-footed mice, forested white-footed mice, and all deer, respectively. Subsequent sequencing revealed infection with divergent betacoronaviruses. The limited SARS-CoV-2 spillback into white-footed mice suggests a low risk of novel SARS-CoV-2 reservoir establishment among and viral variant emergence from this species. However, the detection of non-SARS-CoV-2 divergent betacoronaviruses among white-footed mice highlights the broader, likely underestimated contribution of rodents to betacoronavirus diversity and circulation.

## Results

We investigated whether there was evidence of SARS-CoV-2 spillback into white-footed mice in Connecticut by testing 2020-2022 sera and 2022 swabs (oral and anal) for neutralizing antibodies and active infection, respectively. To provide a comparison, we additionally tested 2021-2022 sera and 2022 swabs (oral, anal, and nasal) collected from white-tailed deer, a known susceptible animal, in similar settings. Finally, we tested swabs from both species for additional coronaviruses via a nested PCR to understand whether non-SARS-CoV-2 coronaviruses were able to circulate within our studied animal populations. We sampled white-footed mice from residential (Guilford, CT) and forested (North Branford, CT) settings and deer from two forested settings surrounded by residential neighborhoods (Norwalk and Bridgeport, CT) as part of ongoing host-targeted tick management studies^36–38^ (**Figure 1A**).

**Figure 1:**
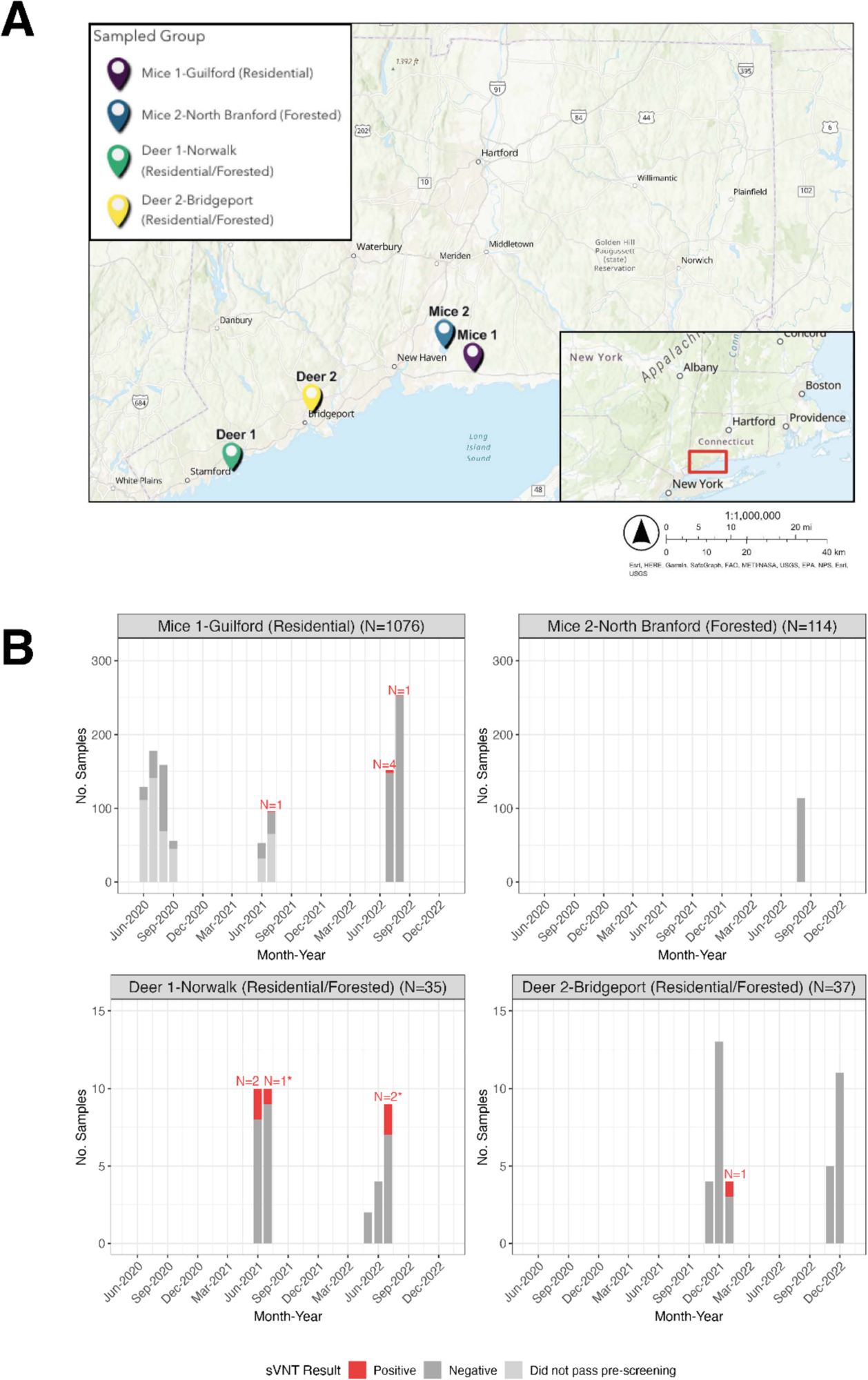
A) Location of study samples in Connecticut. B) Wild-type SARS-CoV-2 neutralizing antibodies detected in white-footed mice and white-tailed deer at various sampling locations in Connecticut. A) Map of sampling locations for white-footed mice in Guilford (Mice 1) and North Branford (Mice 2) and white-tailed deer in Norwalk (Deer 1) and Bridgeport (Deer 2) in Connecticut. The pin location corresponds to the mean coordinates for samples for which we had precise geographic locations (Mice 1). For the other groups (Mice 2, Deer 1, and Deer 2), the pin location shows the approximate sampling area. The inset shows the approximate sampling area within Connecticut. B) Monthly number of sera samples and wild-type neutralizing antibody results for white-footed mice in residential and forested settings and deer in two settings at the residential/forest interface. For the 2020-2021 white-footed mice sera, we only pre-screened the last sample in the case of multiple recaptures, and thus these are not included in the total counts. In addition, we only tested 2020-2021 white-footed mouse samples for neutralizing antibodies if they passed the pre-screening ELISA (**Supplementary Figure 1**). For 2022 white-footed mice and 2021/2022 deer, we forwent the pre-screening ELISA and tested all samples for neutralizing antibodies via the sVNT. We tested the samples either in duplicate initially or in singlicate with a subsequent confirmatory duplicate in the case of an initial positive, unless there was insufficient volume. Note the different y-axes for each plot. *\**Indicates a second positive for an individual recaptured animal. One deer in July 2021 previously tested positive in June 2021. An additional deer tested positive twice during July 2022.

### Wild-type SARS-CoV-2 neutralizing antibodies detected in 1% of residential white-footed mice and 7% of white-tailed deer

To evaluate past SARS-CoV-2 exposure in white-footed mice, we adopted a two-step screening process for sera samples. First, to select 2020 and 2021 white-footed mice with probable past SARS-CoV-2 exposure, we developed an in-house pre-screening ELISA assay to detect binding antibodies (**Supplementary Figure 1**). We did not pre-screen the 2022 white-footed mice samples and 2021/2022 deer samples due to a relatively higher expected prevalence and smaller number of samples. Together with the 2020/2021 samples that passed the pre-screening, we tested sera for neutralizing capacity against wild-type SARS-CoV-2 using the Genscript cPass surrogate virus neutralization test (sVNT). The assay measures neutralizing capacity as the ability to block or reduce the interaction of the viral spike glycoprotein RBD and the human ACE2 receptor pre-coated on the plates^39^.

Among individual residential white-footed mice, we found wild-type SARS-CoV-2 neutralizing antibodies among zero individuals from 2020 (0/156), one from 2021 (1/52, 2%), and five from 2022 (5/332, 2%) (**Figure 1B**). Of the six neutralizing antibody positive samples, one was positive in duplicate on a subsequent confirmatory test (high confidence in positive result), four were positive on the first sVNT test and had elevated OD values that did not surpass the positivity cutoff on the confirmatory test (moderate confidence in a positive result), and one was positive on the first test but was unable to be retested due to insufficient serum volume. Two of the positive white-footed mice were recaptured with discordant results between subsequent samples. One individual initially tested positive for neutralizing antibodies, followed 21 days later by a negative sample.

Another individual initially tested negative for neutralizing antibodies, followed 17 days later by a positive sample. While the sVNT assay is qualitative, all positive white-footed mouse sera samples had values near the assay positivity cutoff, potentially indicating the presence of neutralizing antibodies below the necessary concentration to pass the assay positivity cut-off. Among individual residential white-footed mice, the sex breakdown was as follows: 47% female, 51% male, and 2% unknown. We did not detect any neutralizing antibody positives among the forested white-footed mice (0/69 individual animals). Individual forested mice were 43% female, 54% male, and 3% unknown. The results indicate a limited presence of neutralizing antibody titers against SARS-CoV-2 sufficient to pass the assay positivity threshold, suggesting either absent or infrequent past exposure.

Among the deer sera samples, we identified six positives for wild-type SARS-CoV-2 neutralizing antibodies across 2021 (3/41, 7%) and 2022 (3/31, 10%) (**Figure 1B**). Two of the positives represented recaptured individuals that later tested positive again 16 and 37 days later, respectively, corresponding to four unique individual positives. Only one recaptured deer had discordant serological results possibly indicating waning antibody titers - an initial positive followed 309 days later by a negative sample. Seroprevalence was 12% (3/25) among individual Norwalk deer and 3% (1/30) among individual Bridgeport deer, corresponding to an overall seroprevalence of 7% (4/55). The sex breakdown was as follows: 56% female, 44% male, and 0% unknown at the Norwalk site and 50% female, 47% male, and 3% unknown at the Bridgeport site. The deer were 80% adult, 16% yearling, 4% age transitioning (fawn or yearling to adult in the case of recaptured individuals) at the Norwalk site and 77% adult, 7% yearling, 7% age transitioning, and 10% fawn at the Bridgeport site. Our findings indicate that white-tailed deer in residential/forested settings in Connecticut exhibit higher SARS-CoV-2 neutralizing antibody seroprevalence than white-footed mice sampled from similar habitats, suggesting that the mice potentially are not susceptible or exposed enough in real-world settings for frequent SARS-CoV-2 infection and transmission.

Separately for each species, we conducted a Fisher’s exact test of independence to assess whether there was a significant association between the wild-type SARS-CoV-2 neutralizing antibody result (positive or negative) for each individual and sampling location. We counted each individual animal once. If a recaptured individual ever tested positive, we designated it positive. We failed to reject the null hypothesis for both the white-footed mice and the deer, indicating that there is not a significant association between SARS-CoV-2 neutralizing antibody result and sampling location. In addition, most of the neutralizing antibody positive residential white-footed mice were spatially dispersed, reducing the likelihood of any transmission between mice (**Supplementary Figure 2**). Finally, we also conducted a Fisher’s exact test of independence for both the mice and deer, separately, to test for a significant association between sex (deer and mice) or age (deer only) and neutralizing antibody result. We failed to reject the null hypothesis for all tests, indicating that there is not a significant association between sex or age and neutralizing antibody result.

### Waning or absent cross-neutralization of subsequent SARS-CoV-2 variants observed among white-footed mice and white-tailed deer positive for wild-type SARS-CoV-2 neutralizing antibodies

To understand the possible timing of past exposure and susceptibility to various SARS-CoV-2 lineages, we tested white-footed mice and white-tailed deer samples positive for wild-type SARS-CoV-2 neutralizing antibodies against several variants. Given limited sera volume, we only tested white-footed mice samples in singlicate against the most recent SARS-CoV-2 variant circulating in humans at the time of sample collection^40^. To do so, we used the sVNT as previously described, exchanging the wild-type viral spike glycoprotein RBD for variant-specific RBDs. We tested the July 2021 seropositive white-footed mouse sample against Delta (B.1.617.2) and four samples collected in July/August 2022 against Omicron (BA.2). None of the tested mice sera cross-neutralized RBD variants beyond WT-RBD.

Of the six wild-type neutralizing antibody positive deer samples (including two recaptures), four successfully neutralized Alpha (B.1.1.7) (**Figure 2**). Of those, only one neutralized Delta (B.1.617.2). Following the variant-specific sVNT testing, we further used deer sera to conduct an infectious virus neutralization test (VNT) to confirm our results (**Supplementary Figure 3**) using wild-type, Delta (B.1.617.2), and Omicron (BA.5) viruses. The majority of the deer sera showed neutralizing capacity against wild-type SARS-CoV-2 virus but not against Delta (B.1.617.2) or Omicron (BA.5).

**Figure 2:**
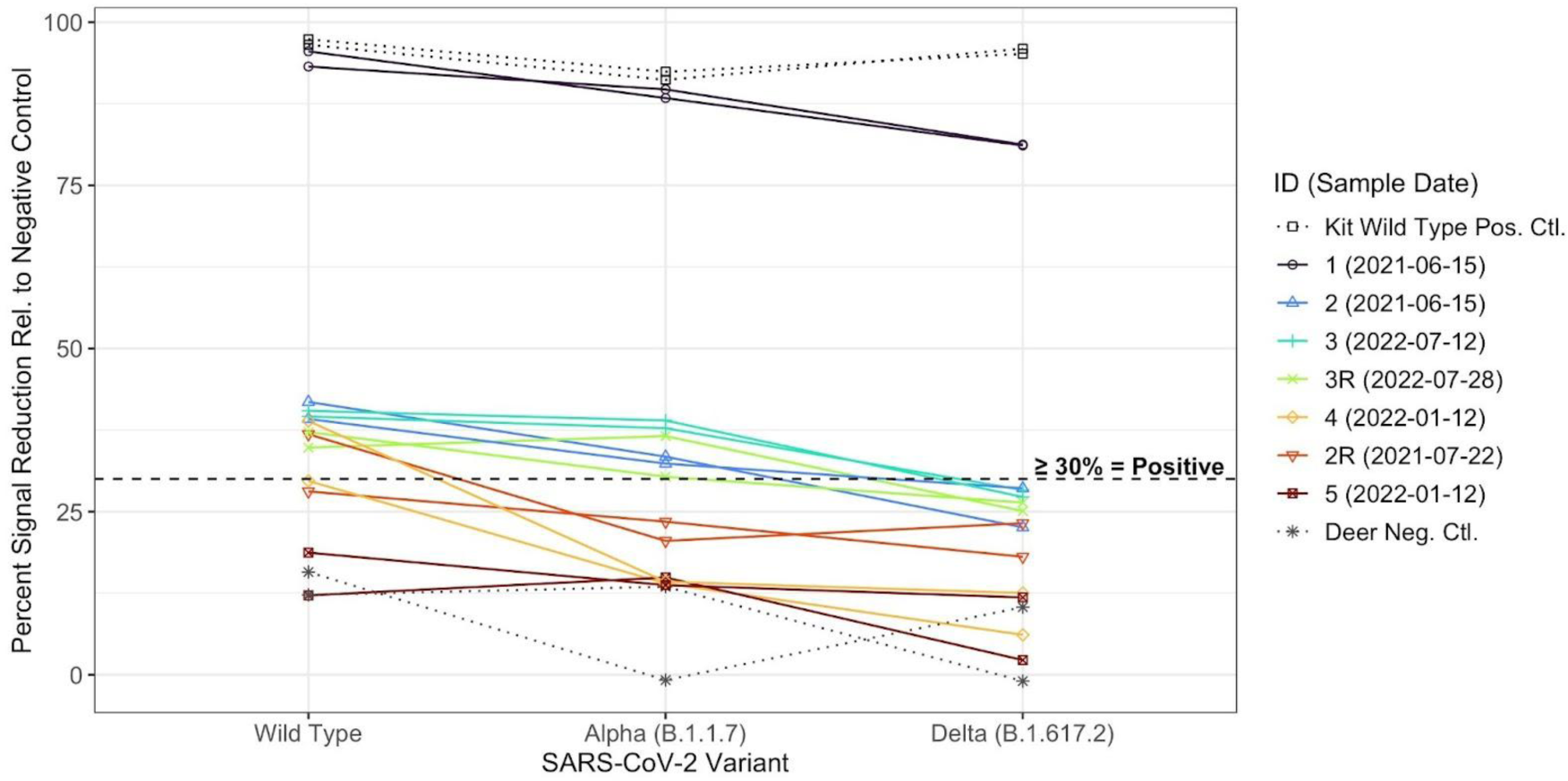
Waning or absent cross-neutralization of Alpha (B.1.1.7) and Delta (B.1.617.2) viral variants among wild-type SARS-CoV-2 neutralizing antibody positive deer. Percent signal inhibition for each SARS-CoV-2 variant relative to the kit negative control using the sVNT. We tested each sample in duplicate at 1:10 dilution against wild-type, Alpha (B.1.1.7), and Delta (B.1.617.2). We included wild-type SARS-CoV-2 kit controls, as well as a wild-type neutralizing antibody negative deer. The dashed line indicates the ≥ 30% positivity cutoff. The sample IDs in the legend are in order of descending mean percent inhibition for wild-type SARS-CoV-2. “R” at the end of a sample ID indicates a recaptured animal.

### No active SARS-CoV-2 infections detected via RT-qPCR among 2022 swabs from white-footed mice and white-tailed deer

We collected two swabs (oral and anal) per mouse and three swabs (oral, anal, and nasal) per deer to test for active SARS-CoV-2 infection (**Supplementary Figure 4**). We sampled white-footed mice from July-August 2022 and white-tailed deer from May-July and November-December 2022, timed to coincide with ongoing studies that enabled our sample collection^36–38^. Most sampling occurred during periods of low to moderate estimated human transmission^41^. We pooled the swab samples for each individual animal on each collection date for RNA extraction and RT-qPCR testing^42^ (**Supplementary Figure 5**). We did not detect any SARS-CoV-2 PCR positives among either the mice (n=614) or the deer (n=31) pooled swabs.

### Pan-coronavirus PCR testing followed by sequencing revealed divergent betacoronavirus infections among 1% of residential white-footed mice sampled in 2022

To understand whether non-SARS-CoV-2 coronaviruses were capable of circulation among our studied animal populations, we tested all 2022 pooled white-footed mice and deer swabs for coronaviruses from four genera (alpha-, beta-, gamma-, and deltacoronaviruses; **Supplementary Figure 5**). We used a semi-nested pan-coronavirus PCR assay that targets a highly conserved region of the RNA-dependent RNA polymerase (RdRp) gene^43^. A recent study performed a comparison of pan-coronavirus PCR approaches using a set of novel bat coronaviruses and found that our selected approach outperformed 3/4 comparison approaches^44^. Among tested pooled swabs collected in 2022, we noted a 1% (5/468), 0% (0/146), and 0% (0/31) pan-coronavirus PCR positivity among the residential white-footed mice, forested white-footed mice, and deer, respectively. We detected two spatiotemporal groupings among our positives: 1) two positive residential mice sampled on the same day from the same residential property and 2) three positive mice sampled on the same day from three residential properties maximally separated by approximately 160 meters (**Figure 3A**). We did not detect a significant association between sex and pan-coronavirus PCR result as measured by a Fisher’s exact test of independence. Our findings point to low-level circulation of non-SARS-CoV-2 coronaviruses among our studied white-footed mice population.

**Figure 3:**
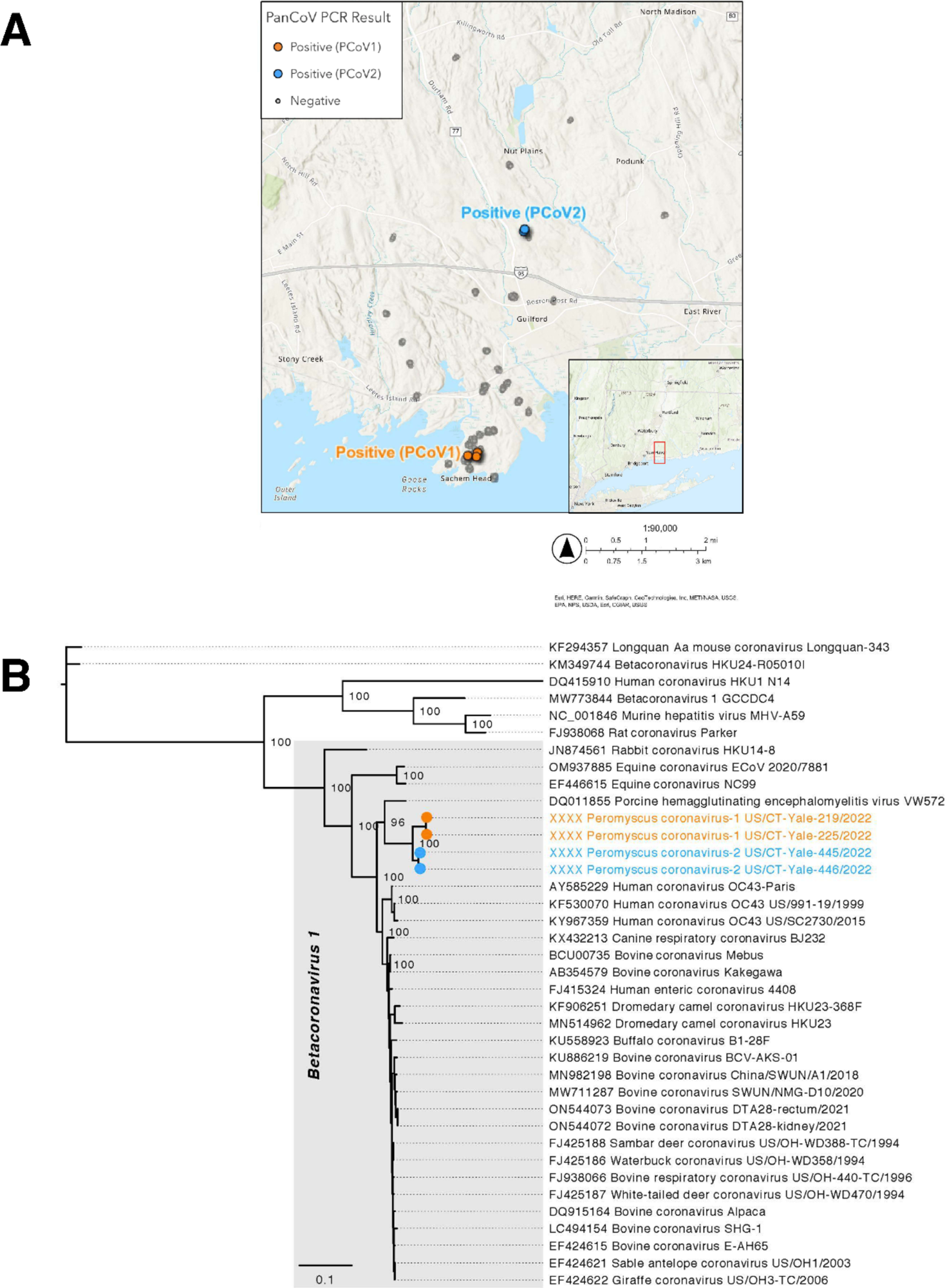
A. Two clusters of pan-coronavirus PCR detections among white-footed mice sampled from a residential setting in Guilford, CT. B) Phylogenetic tree of divergent betacoronavirus genomes recovered from residential white-footed mice. A) The three positive mice in the PCoV1 cluster were sampled on the same day from three residential properties maximally separated by approximately 160 meters, within the potential home range of white-footed mice^46^. The two positive mice in the PCoV2 cluster were sampled on the same day from the same residential property. The colors correspond to the phylogenetic tree shown in **Figure 3B**. We did not display untested samples and jittered the sample location coordinates for visibility. The inset displays the sampling location within Connecticut. B) We obtained four complete genomes from the five pan-coronavirus PCR positive samples in **Figure 3A** via metagenomic amd amplicon sequencing. The sample sequences formed two distinct betacoronavirus clusters: PCoV1 and PCoV2. GenBank accession numbers will be provided when they are available (see Data Availability statement).

We further confirmed the specificity of the PCR product via sequencing and subsequently performed metagenomic sequencing on confirmed positive samples (**Supplementary Figure 5**). We identified divergent betacoronaviruses among the four samples for which we were able to obtain complete viral genomes (**Figure 3B**). We observed two distinct viral clusters that we preliminarily designated *Peromyscus coronavirus-1* (PCoV1) and *Peromyscus coronavirus-2* (PCoV2) (within cluster identity = 99.57-99.96%; between cluster identity = 96.51-96.68%). PCoV1 and PCoV2 are most closely related to *Porcine hemagglutinating encephalomyelitis virus* (PHEV; GenBank accession DQ011855) at 88.68-89.37% similarity, and they are nested within the *Betacoronavirus 1* clade (*Betacoronavirus* genus, *Embecovirus* subgenus^45^; **Figure 3B**). The *Betacoronavirus 1* species includes various human (e.g. *human coronavirus OC43*) and animal coronaviruses (e.g., *bovine coronavirus*, *equine coronavirus*, *canine respiratory virus*). In **Supplementary Figure 6**, we report the sequencing coverage and display phylogenetic trees for the ORF1ab, spike, and nucleocapsid genes. We found the greatest genetic divergence in the spike. PCoV1 and PCoV2 are 99.05-99.09%, 89.93-90.08%, and 95.03-95.10% identical to each other in the ORF1ab, spike, and nucleocapsid genes, respectively. PCoV1 and PCoV2 are 92.20-92.36%, 85.41-86.97%, and 92.89-94.07% identical to PHEV in the ORF1ab, spike, and nucleocapsid genes, respectively. Additional sequencing and GenBank submission details are available in **Supplementary Tables 1-2.**

## Discussion

To understand the long-term SARS-CoV-2 risk posed by white-footed mice in Connecticut, we tested sera for wild-type neutralizing antibodies via sVNT, finding 1% and 0% seroprevalence among individual white-footed mice in the residential and forested settings, respectively (**Figure 1B**). To provide a comparison, we assesed seroprevalence among white-tailed deer, a species with high levels of reported exposure and infection, from similar Connecticut settings. We observed 7% neutralizing antibody seroprevalence across the two residential/forested settings. We subsequently tested wild-type neutralizing antibody positive mice sera from 2021 and 2022 against Delta (B.1.617.2) and Omicron (BA.2), respectively, observing no cross-neutralization. We tested neutralizing antibody positive deer sera against infectious Alpha (B.1.1.7), Delta (B.1.617.2), and Omicron (BA.5), noting waning or absent cross-neutralization (**Figure 2 and Supplementary Figure 3**). We did not identify active SARS-CoV-2 infections via RT-qPCR in swabs from either species (**Supplementary Figure 4**). Lastly, to understand broader coronavirus circulation, we performed pan-coronavirus PCR testing of all 2022 mice and deer swabs and noted 1% (5/468), 0% (0/146), and 0% (0/31) pan-coronavirus PCR positivity among the residential white-footed mice, forested mice, and deer, respectively (**Figure 3A**). Sequencing yielded complete genomes from four of the five positive mice samples, revealing two divergent viruses (preliminarily named PCoV-1 and -2) that cluster with members of the *Betacoronavirus 1* species (**Figure 3B and Supplementary Figure 6**). Our findings indicate low spillback of SARS-CoV-2 from humans or an unknown animal into white-footed mice, with sustained transmission and secondary spillover unlikely. However, our detection of divergent betacoronaviruses in white-footed mice demonstrates that non-SARS-CoV-2 coronaviruses are capable of circulation. Further research is required to understand the extent of infection with the newly identified betacoronaviruses.

Among the 1% of residential white-footed mice samples that initially tested positive for wild-type SARS-CoV-2 neutralizing antibodies (**Figure 1B**), several displayed elevated values that did not cross the 30% positivity threshold upon retesting. First, neutralizing antibodies could be present but at insufficient levels to consistently surpass the assay cutoff. Second, there may have been false positives on the initial test. Third, individuals may not have been exposed to SARS-CoV-2, but rather have cross-reactive antibodies to an unknown coronavirus (possibly PCoV). Finally, because we confirmed the initial positive test with a subsequent confirmatory test, the difference could reflect slight variation in how the assay was conducted. One mouse positive for SARS-CoV-2 neutralizing antibodies was recaptured 21 days later and tested negative. In addition to the previously mentioned hypotheses for differential sVNT results, this finding could reflect seroconversion in the interim period between sample collections or diminished antibodies following the initial blood draw. We did not detect any variant cross-neutralization among tested sera from wild-type seropositive mice. Our finding may be due to exposure to an earlier SARS-CoV-2 lineage combined with low cross-neutralization between variants^47^. The white-footed mice population typically turns over annually due to high mortality as demonstrated by the few interannual recaptures in our study^46^. Therefore, our SARS-CoV-2 neutralizing antibody findings would maximally include past infections dating back approximately one year, if neutralizing antibodies persist for that duration. White-footed mice also experience the highest levels of both reproduction and mortality during summer and the opposite dynamics during winter^48,49^. Thus, our summer sampling may have captured relatively younger mice with potentially lower overall seroprevalence. Of the neutralizing antibody positive mice, only two were captured within potentially overlapping territories (**Supplementary Figure 2**)^46^. The low SARS-CoV-2 neutralizing antibody seroprevalence combined with the spatial dispersion of positives does not suggest sustained transmission between mice following occasional spillbacks.

While the 7% wild-type SARS-CoV-2 neutralizing antibody seroprevalence observed among white-tailed deer in our study (**Figure 1B**) was lower than the 14-40% neutralizing antibody positivity found in other locations^10,23,28,30,32^, it was surprisingly high given the semi-restricted nature of the sampling sites. All six seropositive deer samples tested positive in duplicate. One seropositive deer tested negative upon recapture 309 days later, likely due to waning immunity. We tested wild-type seropositive deer against Alpha (B.1.1.7) and Delta (B.1.617.2), observing absent or decreasing neutralization across variants (**Figure 2**). Four samples successfully neutralized Alpha and, among those, one neutralized Delta. We observed the highest percent inhibition of wild-type SARS-CoV-2, suggesting past infection with wild-type SARS-CoV-2. Seropositive deer may have been exposed in 2020 prior to the spread of variants in deer populations or may have been more recently exposed to an earlier lineage potentially still in circulation in the animal population. Regardless, we cannot rule out subsequent variant infections if the deer mounted the strongest neutralizing antibody response to their initial infection^50^. Deer #1 displayed especially high neutralizing activity against wild-type, Alpha, and Delta, possibly due to a more recent infection or a stronger immune response to infection due to individual immunological variation. Deer #5 may have been a false positive on the initial wild-type SARS-CoV-2 sVNT. We subsequently tested sVNT positive deer using an infectious VNT (**Supplementary Figure 3**). Most deer neutralized wild-type SARS-CoV-2 and none neutralized Delta or Omicron (BA.5). Assay result variation could be due to differential ACE2 receptor presentation, the use of only the viral spike protein RBD in the sVNT versus infectious virus in the VNT, or that the sVNT measures inhibition of ACE2/viral spike protein RBD binding whereas the VNT measures cell viability. White-tailed deer live up to 10 years in the wild^51^ and one study indicated that neutralizing antibodies persist for at least 13 months in the majority of naturally-infected deer^52^. Thus, as all of the seropositive deer in our study were adults excluding one yearling (#1), their infections could potentially date back to at least 13 months prior to sampling. Deer home ranges vary from 0.8-3.2 and 1.6-6.4 square kilometers for adult females/fawns and adult males, respectively^51^. Adult males substantially increase their movements during the late fall breeding season and yearling males can disperse up to 32 kilometers to establish new home ranges. The higher wild-type SARS-CoV-2 neutralizing antibody seroprevalence observed within white-tailed deer compared to other species could in part result from such movement patterns, with infections in new herds seeded by adult or yearling males.

None of the mice or deer samples tested positive via RT-qPCR for SARS-CoV-2 infection, potentially due to the relatively low-risk sampling periods timed to coincide with ongoing studies^36–38^ (**Supplementary Figure 4**). We primarily collected samples during summer 2022, which corresponded to a period of lower human transmission and lower-risk animal behavior. Both species are relatively solitary during summer but form larger groups during winter^53–55^. In addition, mice experience greater human exposure during winter when they enter human homes seeking warmth. However, we also did not detect any infections among deer sampled during the relatively higher-risk colder months. In addition, the window of active infection detection may be relatively narrow. Experimental SARS-CoV-2 challenge studies of *Peromyscine* mice found detectable viral RNA and infectious virus up to 8-21 and 3-14 days post infection, respectively, (ranges across studies) in oral swabs and at least eight days in rectal swabs^13–16^. Challenged deer had detectable viral RNA up to 7-14, 10-22, and 7 to at least 14 days post infection in oral, nasal, and rectal swabs, respectively (ranges across studies) and infectious virus from all swab types up to five days^33–35^. However, challenge studies typically use higher viral doses than those naturally encountered, and thus real-world detection windows may be narrower. Finally, our sampling locations likely feature a low risk of SARS-CoV-2 exposure from humans. We sampled residential mice from the backyards of single-family homes in a relatively affluent suburban area. In contrast, mice residing in or around high-density or poorer quality housing that enables easier access would likely experience greater exposure. Similarly, deer sampled from locations with greater access to human populations or other deer herds would likely have a higher exposure risk.

We detected 1% pan-coronavirus PCR positivity among residential white-footed mice (5/468) (**Figure 3A**) and sequenced complete viral genomes from four samples (**Figure 3B**). We detected two divergent betacoronavirus clusters (PCoV1 and PCoV; 96.51-96.68% identical) that aligned spatiotemporally with sample collection. Given such divergence between geographic clusters combined with the limited range of *Peromyscine* mice, transmission of PCoV1 and PCoV2 is likely geographically isolated. Comparing PCoV1 and PCoV2, our phylogenetic trees for specific gene sequences (**Supplementary Figure 6**) revealed nearly identical genomes for the ORF1ab gene (99.05-99.09% identical) with greater divergence in the spike gene (89.93-90.08% identical). PCoV1 and PCoV2 were comparatively more similar in the nucleocapsid gene (95.03-95.10% identical), but PCoV1 appeared to be ancestral to PCoV2, potentially a result of recombination. The PCoV1 and PCoV2 sequences clustered most closely with *Porcine hemagglutinating encephalomyelitis virus* (PHEV; 88.68-89.37% identical), a common virus found worldwide in pigs^56^. However, PCoV1 and PCoV2 remain relatively divergent from PHEV, suggesting that the viral emergence event in white-footed mice (or in another animal species) occurred in the distant past. PCoV1 and PCoV2 both belong to the *Betacoronavirus 1* species (*Betacoronavirus* genus, *Embecovirus* subgenus)^45^. Based on available GenBank sequences with host and location data, prior *Betacoronavirus 1* detections in rodents are limited to human coronavirus OC43 among hamsters (*Cricetinae sp.*) and bovine coronavirus among Daurian ground squirrels (*Spermophilus dauricus*) in China^57,58^. However, the broader *Embecovirus* subgenus has become predominantly associated with rodent hosts in recent years, with detections increasing from one viral species in 2015 (*murine coronavirus*) to at least four additional viral species as of 2023 (*Betacoronavirus 1*, *Myodes coronavirus*, *China Rattus coronavirus HKU24*, and unclassified)^45,57,59,60^. In North America, only the *murine coronavirus* species (*Embecovirus* subgenus) has been reported in rodents^58^. Despite recent advancements, our understanding of rodent coronaviruses remains limited and merits further exploration, particularly given the limited host restriction observed among rodent embecoviruses and hypothesized origin of *human coronavirus OC43* in rodents^57,59^

We potentially underestimated SARS-CoV-2 exposure due to insufficient neutralizing antibodies for detection despite past exposure. The sVNT assay has moderate to high sensitivity (91-94%) and high specificity (100%), but is less suitable for detecting low titers^61^. Our ELISA and sVNT assays detected wild-type SARS-CoV-2 antibodies and animals infected with subsequent variants may test negative. We selected the sVNT assay for its species-agnostic aspect and appropriateness for biosafety level 2 (BSL-2) settings. However, the sVNT assay uses only the viral spike protein RBD and other more conserved parts of the Spike protein may also serve as neutralizing antibody targets. The viral spike protein RBD, while under strong evolutionary pressure, does not reflect all viral genome-wide mutations that differentiate SARS-CoV-2 variants, and there may be greater differences between variants in VNTs using infectious virus that could explain discrepancy in results between assays. In addition, the assay uses human ACE2 receptors. Cross-reactivity to other non-SARS-CoV-2 coronaviruses could explain some of the elevated results that oscillated near the sVNT positivity threshold. The sVNT assay manufacturer reports cross-reactivity with SARS-CoV-1 and MERS, but not with common human coronaviruses^62^. Various alpha- and non-SARS-CoV-2 betacoronaviruses have been recently detected in wild rodent populations^57,59,63–69^, including those identified in our study. In addition, bovine betacoronaviruses have been detected in wild white-tailed deer and other ruminants^70^.

In conclusion, our combined serological and RT-qPCR results suggest limited SARS-CoV-2 spillback from humans or an unknown animal species into white-footed mice at sampled sites in Connecticut, although the absence of active SARS-CoV-2 infections precludes more detailed viral genomic analysis. The ubiquity of the ACE2 receptor among vertebrates combined with the unprecedented exposure opportunity of the ongoing COVID-19 pandemic has provided innumerable SARS-CoV-2 spillback opportunities into naive animal species. As the virus is introduced into a greater range of animal species, there is further potential for spillback between animal species as well. In addition, our detection of divergent betacoronavirus infections among white-footed mice expands the diversity of coronaviruses discovered among rodents. Further investigation is necessary to determine the geographic and species range of PCoV1 and PCoV2. Given the relatively small territories of white-footed mice, it is possible that such viruses may remain constrained to narrow geographic areas unless they circulate among additional animal species with greater movement ranges. Deeper surveillance of animal populations with the susceptibility and behavior potential to serve as potential primary or secondary betacoronavirus reservoirs is key to understanding the possible implications for long-term control of both established and newly detected betacoronaviruses.

## Methods

### Ethics Statements

We followed animal capture and handling protocols approved by the Wildlife Division of the Connecticut Department of Energy and Environmental Protection (#2124002, 2124002a, and 2023006) and The Connecticut Agricultural Experiment Station’s (CAES) Institutional Animal Care and Use Committee (IACUC) (P32-20 and P35-21) in accordance with the American Society of Mammologist’s guidelines for the use of wild animals in research^71^. The Yale University IACUC determined that use of remnant samples from CAES research did not necessitate additional oversight.

### Sample Collection

#### White-Footed Mice Sample Collection

We sampled white-footed mice from a residential area in Guilford, CT and a forested area in North Branford, CT from June-August 2020-2022 as part of ongoing host-targeted tick management studies investigating the effectiveness of systemic fipronil treatments via baited feed boxes^36,37^ (**Figure 1A**). We collected sera samples during all years, with additional oral and anal swabs only in 2022. The white-footed mice were trapped using Sherman live animal traps (LFAHD folding trap, H. B. Sherman Traps, Inc.) baited with peanut butter. In the residential area, 12 traps were set at each of 50 different properties 10 meters apart (on multiple occasions) along the periphery of the lawn/woodland edge. In the forested area, we set traps at 5 locations in a 6×6 grid 10 meters apart six times in 2021 and four times in 2022. Regardless of location, traps were placed in the late afternoon and collected the subsequent morning. Any non-target animals were released. For sample collection, each mouse was transferred to a plastic bag with a cotton ball soaked in a small quantity of the inhalant anesthetic isoflurane (Piramal Critical Care, Inc.) for temporary sedation. Once the animal was sedated, a study team member drew 50-150 μl of blood (<10% of total blood volume) via cardiac puncture contingent upon the animal’s weight, attached a unique metal ear tag (#1005-1, National Band and Tag Co.) to enable identification of recaptures, and collected oral and anal swabs (Puritan 6” Sterile Mini-tip Polyester Swab w/Ultra-Fine Polystyrene Handle 25-800 1PD 50)^72^. Swabs were placed in screw-cap tubes containing viral transport media (VTM) (96% DMEM, 2% FBS, and 2% antibiotic-antimycotic at final concentrations) (Gibco). We did not take blood samples from the same mouse more than once every two weeks. White-footed mice were returned to their Sherman trap until alert and then released to their original collection site. Sera and swabs were stored at -80°C.

#### White-Tailed Deer Sample Collection

We sampled deer from two forested areas with abutting urban development at least partially separated by a chain-link fence in Norwalk and Bridgeport, CT from May-July and November-January in 2021 and 2022 (**Figure 1A**). We collected sera each year as part of a tick management study investigating systemic moxidectin treatment via timed feeding stations^38^, with additional oral, nasal, and anal swabs only collected in 2022. At designated sites in both locations, the study team set up timed feeders that released food near dusk. Trained staff used transmitter darts with a combination of 2.0 ml butorphanol tartrate, azaperone tartrate, and medetomidine hydrochloride (BAM). Radio telemetry equipment was used to locate the dart to determine the sedated animal’s location. A study team member drew 6.0 ml of whole blood via venipuncture from each deer. Oral, anal, and nasal swabs were also taken (Puritan 6” Sterile Standard Polyester Swab w/ Polystyrene Handle REF 25-806 1PD for oral and anal swabs; LIBO Specimen Collection and Transport Swabs Item No 30151 for nasal swabs) and placed into snap-lock or screw-cap tubes with VTM. We added ear tags to each animal to enable reidentification in the case of recapture (All-Flex). After sample collection, the effects of BAM were antagonized with a combination of 3.0 ml atipamezole and 0.5 ml naltrexone delivered via intramuscular injection. Sera and swabs were stored at -80°C.

### Serological Testing, PCR Testing, and Sequencing

#### Sera Sample Inactivation

We inactivated sera samples using either detergent or heat in a BSL-2 setting. We treated detergent-inactivated samples with a solution of Triton X-100 10% (0.5% final concentration) (Sigma Aldrich) and RNase-A 100 mg/mL (0.5 mg/mL final concentration) (Qiagen). We treated heat-inactivated samples at 56°C for 30 minutes.

#### ELISA

Due to low expected SARS-CoV-2 exposure prevalence and a large volume of samples, we used an in-house ELISA assay to pre-screen all 2020/2021 white-footed mice sera samples for IgG binding antibodies (**Supplementary Figure 1**). We only pre-screened the last sample in the case of multiple recaptures. Samples with OD values above a cutoff threshold were tested for wild-type SARS-CoV-2 neutralizing antibodies. We directly tested the 2022 white-footed mice and all deer sera samples for neutralizing antibodies without the pre-screening ELISA.

We began by pre-screening the 2021 samples (plates 1-4) (**Supplementary Figure 7**), hypothesizing that these would be more likely to include a positive mouse than the 2020 samples. At this time, we had not initiated sera collection for the 2022 white-footed mice. For plate 1, given our initial lack of a positive white-footed mouse control, we tested each sample in triplicate on wells separately coated with SARS-CoV-2 spike, SARS-CoV-2 nucleocapsid, and BSA proteins. We found similar values for the nucleocapsid and BSA proteins and thus dropped the nucleocapsid protein coating to streamline sample processing. We used 2010 white-footed mice samples as the negative control. Any samples with a spike protein value ≥1.57x (a convenience cutoff) the BSA protein value were tested for neutralizing antibodies. After detecting one neutralizing antibody positive mouse among the 2021 samples, we included it as the positive control for the 2020 sample pre-screening (plates 5-16), stopped the BSA protein comparison, and re-added the nucleocapsid protein coating. All remaining samples were tested simultaneously on spike and nucleocapsid protein coated plates. After plates 5/6, we also removed the 2010 white-footed mice negative controls as they displayed unusually elevated values compared to the positive control, possibly due to the sample age. Instead, we used both a neutralizing antibody negative 2020 mouse sample and buffer only as negative controls. Given the low expected exposure among the white-footed mice, we used a conservative positivity cutoff: sample OD values for both the spike and nucleocapsid protein plates ≥1x the corresponding positive control values.

All materials were from an ELISA Buffer Kit (Invitrogen CNB0011) unless otherwise indicated. We diluted the samples 1:500 with a blocking buffer. We diluted the spike (ECD, His & Flag Tag) (Genscript <u>Z03481</u>), nucleocapsid (Genscript Z03488), and BSA proteins to 0.5 ug/mL concentration using carbonate buffer. For the detection antibody, we diluted 1 mg/mL Anti-Peromyscus Leucopus IgG (H+L) Antibody Peroxidase-Labeled (Seracare 5220-0375) 1:10K using a blocking buffer. For each run, we coated two 96-well flat-bottom plates (Thermo Scientific Clear Flat-Bottom Immuno Nonsterile 96-Well Plates 439454) with spike and nucleocapsid protein, separately, and incubated them at 4°C overnight. The following day, we washed the plates four times with 1X Wash Buffer, added a blocking buffer, incubated the plates for one hour at room temperature, and washed the plates four times. We added samples in singlicate to each plate. We then incubated the plates for two hours at room temperature, washed the plates four times, added the detection reagent, incubated the plates for one hour at room temperature, and washed the plates four times with 1X Wash Buffer followed by one wash with 1X PBS. Finally, we added stabilized chromagen and quenched the reaction after ∼15 minutes with a stop solution. The plates were immediately read at OD 450 nm on a plate reader.

#### Surrogate Virus Neutralization Test

We used a surrogate virus neutralization test (sVNT) (Genscript cPass Ref L00847) to detect wild-type SARS-CoV-2 neutralizing antibodies (**Figure 1B**)^39^. The assay detects neutralizing antibodies that block the interaction of wild-type SARS-CoV-2 spike protein RBD with ACE2 receptor pre-coated on the plate. We used kit negative and positive controls, each in duplicate. We tested the samples either in duplicate initially or in singlicate with a subsequent confirmatory duplicate in the case of an initial positive.

We generated neutralization reaction mixtures by combining the samples with 1:1K diluted RBD-HRP solution on a prep plate (Thermo Fisher Clear Round-Bottom Immuno Nonsterile 96-Well Plates MaxiSorp 449824). We incubated the prep plate for 30 minutes at 37°C. Next, we added the neutralization reaction mixtures to the pre-coated capture plate and incubated it for 15 minutes at 37°C. We washed the capture plate four times with 1X Wash Solution, added TMB solution, incubated the plate in a drawer for 15 minutes at ∼25°C, and added stop solution to quench the reactions. The plates were immediately read at OD 450 nm on a plate reader. We calculated the percent signal inhibition relative to the mean (across duplicates) kit negative controls, and used the manufacturer-recommended ≥ 30% signal inhibition positivity cutoff. To assess cross-neutralization using the same sVNT assay, we exchanged the default wild-type viral spike protein RBD for variant-specific RBDs including Alpha (B.1.1.7) (Genscript Z03595), Delta (B.1.617.2) (Genscript Z03614), and Omicron (BA.2) (Genscript Z03741) (**Figure 2**). We used the same positivity cutoff and the kit wild-type SARS-CoV-2 positive control for all variants excluding Omicron (BA.2), for which we used sera from a recently Omicron (BA.5)-infected lab mouse (sera gift of Tianyang Mao) due to the variant’s immune escape properties.

#### Infectious Virus Production and Characterization

To generate viral stock, Vero-E6 ACE2 TMPRSS2+ cells were infected with P3 stock made in Huh7.5 cells using SARS-CoV-2 isolate USA-WA1/2020 (BEI Resources #NR-52281) as previously described^73^. Full-length Delta (B.1.617.2) production and characterization were described previously^74^. Vero-E6 overexpressing ACE2 and TMPRSS2 (VeroE6-AT) were inoculated with SARS-CoV-2 Omicron BA.5 isolate as previously characterized^73,74^.

#### Infectious Virus Neutralization Test

Deer sera and pooled human sera (BEI# NRH-21762) were heat inactivated prior to the infectious virus neutralization assay (**Supplementary Figure 3**). The convalescent human sera was used as a positive neutralization control. Sera samples were serially diluted in serum-free DMEM, then incubated with infectious virus for one hour at 37°C. Samples were tested in parallel by two different technicians, with each plate containing two technical replicates per sample. After coincubation, the sera/virus mixture was added to the VeroE6 ACE2-TMPRSS2 cells at final virus MOI=2. Cell viability was measured at 3 dpi using the CellTiter-Glo® Assay (Promega) on a Cytation 5 (Biotek) luminometer.

#### Nucleic Acid Extraction and Pooled RT-qPCR Testing

We pooled 100 μL from the oral, anal, and nasal swabs collected from each deer and 150 μL from the oral and anal swabs collected from each white-footed mouse (**Supplementary Figure 4**). We extracted nucleic acid from 300 μL of pooled original sample using the MagMAX Viral/Pathogen Nucleic Acid Isolation Kit (ThermoFisher Scientific A42352) on the ThermoFisher KingFisher Flex Purification System, eluting in 75 μl of elution buffer. We tested the extracted nucleic acid for SARS-CoV-2 RNA using a “research use only” (RUO) RT-qPCR assay on the Bio-Rad CFX96 Touch Real-Time PCR Detection System^42^. We included negative extraction and template controls (nuclease-free water) and synthetic SARS-CoV-2 RNA (1000 copies/μl) positive controls^42^.

#### Pan-Coronavirus PCR Testing

We first synthesized cDNA from the nucleic acid extracted during SARS-CoV-2 RT-qPCR testing using 4 μl of SuperScript IV VILO Master Mix (ThermoFisher Scientific 11756050), 6 μl of nuclease-free water, and 10 μl of nucleic acid per reaction (**Figure 3A and Supplementary Figure 5**). We included a positive feline coronavirus control (gift from Cornell University College of Veterinary Medicine) and negative extraction and template controls (nuclease-free water). To test for coronavirus RNA, we followed a semi-nested approach that targets conserved regions of the RNA-dependent RNA polymerase (RdRp) gene and used the accompanying primers as previously described^43^. We also used the primers designed for this approach. We combined 2 μl of each primer (IDT), 25 μl of Platinum II Hot-Start Green PCR Master Mix 2X (Invitrogen 14001014), 15 μl of water, and 4 μl of cDNA per reaction. For the first round, we used the following primers: Pan_CoV_F1 (GGTTGGGAYTAYCCHAARTGYGA), Pan_CoV_R1 (CCRTCATCAGAHARWATCAT), and PanCoV_R2 (CCRTCATCACTHARWATCAT) (IDT). We performed the following thermal cycling steps: 3 minutes at 94°C, 25 cycles of 30 seconds at 94°C, 30 seconds at 48°C, 1 minute at 72°C, and 5 minutes at 72°C. For the second round, we used 2 μl of the first-round PCR product, 2 μl of each primer (IDT), 25 μl of Platinum II Hot-Start Green PCR Master Mix (2X) (Invitrogen 14001014), and 15 μl of water per reaction. We used the following primers: Pan_CoV_R1 (CCRTCATCAGAHARWATCAT), PanCoV_R2 (CCRTCATCACTHARWATCAT), PanCoV_F2 (GAYTAYCCHAARTGTGAYAGA), and PanCoV_F3 (GAYTAYCCHAARTGTGAYMGH) (IDT). We conducted the following thermal cycling steps: 3 minutes at 94°C, 40 cycles of 30 seconds at 94°C, 30 seconds at 58°C, 1 minute at 72°C, and 5 minutes at 72°C. We visualized PCR products on a 2% agarose gel. We initially validated our approach using positive human SARS-CoV-2 (a betacoronavirus) controls and a positive feline coronavirus (an alphacoronavirus) control, using the latter as the positive control for all plates.

#### Sequencing

To confirm the specificity of the PCR products from the panCoV PCR we prepared the PCR products for sequencing on the Illumina NovaSeq 6000 (paired-end 150bp) using the the tagmentation and adapter ligation reaction from the Nextera XT DNA Library preparation kit followed by adding index adapters via PCR for sample identification prior to pooling. For samples where we confirmed PCR product specificity, we subsequently performed untargeted metagenomic sequencing as previously described^75^. Briefly, we re-extracted RNA from each anatomical site separately. Nucleic acid extraction was treated with DNase followed by first- and second-strand cDNA synthesis. The cDNA was directly submitted to the tagmentation and adapter ligation reaction using the Nextera XT DNA Library preparation kit. Then, index adapters were added via PCR for sample identification prior to pooling. Sequencing was performed on a Illumina NovaSeq 6000 (paired-end 150bp) at the Yale Center for Genome Analysis, targeting 50-100 million reads per individual library. The sequencing data are available at NCBI Bioproject PRJNA1003876 (www.ncbi.nlm.nih.gov/bioproject/PRJNA1003876).

### Data Analysis

All data analysis was performed using R (version 4.0.5) and RStudio (version 1.4.1106)^76,77^.

#### Seroprevalence

We calculated the seroprevalence as the number of unique positive individuals divided by the total number of unique tested individuals, unless otherwise noted, as recaptured individuals may repeatedly test positive in the case of recaptures. If a recaptured animal ever tested positive, we counted it as positive.

#### Percent PCR Positivity

We calculated the percent SARS-COV-2 RT-qPCR positive as the number of positive pooled samples (per individual per collection day) divided by the total number of pooled samples.

#### Fisher’s Exact Test of Independence

We used Fisher’s exact test in the R package *stats* (version 4.0.5)^76^ to assess whether there was a significant association between the sVNT wild-type SARS-CoV-2 neutralizing antibody result (positive or negative) for each individual and the sampling locations for each species. The null hypothesis of the test states that the neutralizing antibody result and sampling location are independent; the alternative hypothesis that they are not independent. We selected this test due to fewer than five observations in one category for each species. We compared 1) the residential (Guilford, CT) versus forested (North Branford, CT) setting for the white-footed mice and 2) the two deer settings (Norwalk and Bridgeport, CT). Specifically, we compare counts of negative versus positive individuals in each location. In the case of multiple recaptures, we counted each individual once. If an individual ever tested positive, we designated it as such. In addition, we conducted a Fisher’s exact test to test for a significant association between sex or age, separately, and neutralizing antibody result (positive or negative). We also tested for an association between sex or age and the pan-coronavirus PCR result. Sex data were available for both the mice and deer. We categorized any sex metadata mismatches for recaptured individuals as “Unknown”. Age data were only available for deer (adult, yearling, or fawn). We categorized any recaptured deer that transitioned from one age group to adulthood during the study (i.e. fawn or yearling to adult) as “age transitioned”. We used a significance level of 0.05 for all tests.

#### Estimated Weekly Human Infections per 100,000 Population in Connecticut

We obtained weekly estimated human infections for Connecticut from *Covidestim*, a Bayesian nowcasting model that adjusts for diagnostic and reporting biases by anchoring to more reliable death data (**Supplementary Figure 3**)^41^. We obtained the estimated 2022 state population from the United States Census Bureau to calculate infections per 100,000 population^41,78^.

#### Mapping

We used the R package ggmap (version 3.0.0) to geocode the residential white-footed mice sample locations and ArcGIS Online to create all maps (**Figure 1A**, **Figure 3A, and Supplementary Figure 2**)^79^.

#### Sequencing and Phylogenetic Data Analysis

We performed metagenomic sequencing of pan-coronavirus positives. We performed initial bioinformatic analysis of the sequencing data using Chan Zuckerberg ID (CZ ID, formerly IDseq)^80–82^. We uploaded the raw sequencing reads to CZ ID, which performs host filtering and quality control steps prior to de novo and reference-guided assemblies. We then downloaded all of the contigs that aligned to betacoronaviruses and performed a de novo assembly using Geneious to generate a draft consensus genome. We next generated final consensus genomes for each detected coronavirus by performing a reference-guided alignment of the original sequencing data (fastq files) to the draft consensus genomes using Bowtie 1.1.2 (minimum depth = 10 nucleotides)^83^.

To perform the phylogenetic analysis, we downloaded 34 representative betacoronaviruses from GenBank (listed in **Figure 3B**) based on a previous phylogenetic analysis^57^. We performed a multiple sequence alignment using MAFFT v7.490^84,85^ and constructed a maximum-likelihood phylogenetic tree using PhyML 3.3.20180621 with a GTR substitution model and 100 bootstrap replicates^86^. These steps were repeated for gene-specific analysis shown in **Supplementary Figure 6**. The resulting trees were annotated using FigTree v1.4.2^87^.

## Data Availability

The sequencing data are available at NCBI Bioproject PRJNA1003876 (www.ncbi.nlm.nih.gov/bioproject/PRJNA1003876). The analysis code and alignment and tree files are available on github (github.com/grubaughlab/Peromyscus-CoV). Sample metadata are available upon request. GenBank accession numbers will be provided when they are available.

## Author Contributions

Conceptualization: R.E., A.M.H., S.C.W., N.D.G.; Methods Development: R.E., A.M.H., N.M.F., M.I.B., C.B.F.V., V.E.P., X.A.O.C., L.B.G., C.B.W., M.A.L., S.C.W., N.D.G.; Sample Collection and Processing: R.E., A.M.H., N.M.F., N.F.G.C., R.T.K., A.J.P., A.S., A.G., C.K., H.L., H.R.S., J.L.C., M.A.L., S.C.W., N.D.G.; Laboratory Testing and Analysis: R.E., A.M.H., N.M.F., M.B., R.B.F., Z.Z., M.I.B.; Data Analysis and Interpretation: R.E., A.M.H., N.M.F., R.B.F., V.E.P., N.D.G.; Supervision: C.B.F.V., M.A.L., S.C.W., N.D.G.; Writing - Original Draft: R.E., A.M.H., N.M.F., N.D.G.; Writing - Review & Editing: All authors

## Acknowledgements

We thank B. Menasche for the propagation and characterization of the viral stocks, C. Lucas for providing the SARS-CoV-2 Omicron BA.5 isolate, B. Graham for providing the VeroE6-AT cells, V. Ezenwa, N. Bharti, and D. Weinberger for providing comments on the manuscripts, and P. Jack and S. Taylor for technical advice. This publication was made possible by CTSA Grant Number TL1 TR001864 from the National Center for Advancing Translational Science (NCATS), a component of the National Institutes of Health (NIH) awarded to RE. Its contents are solely the responsibility of the authors and do not necessarily represent the official views of NIH.

## Notes

### Competing Interest Statement

The authors have declared no competing interest.

### Summary of Updates

Figures 1 and 3 were not displaying correctly on some browsers - reformatted to fix issue.

## References

1. Pekar, J. E. et al. The molecular epidemiology of multiple zoonotic origins of SARS-CoV-2. Science 377, 960–966 (2022).

2. Fagre, A. C. et al. Assessing the risk of human-to-wildlife pathogen transmission for conservation and public health. Ecol. Lett. 25, 1534–1549 (2022).

3. Banerjee, A., Mossman, K. & Baker, M. L. Zooanthroponotic potential of SARS-CoV-2 and implications of reintroduction into human populations. Cell Host Microbe 29, 160 (2021).

4. Damas, J. et al. Broad host range of SARS-CoV-2 predicted by comparative and structural analysis of ACE2 in vertebrates. Proc. Natl. Acad. Sci. U. S. A. 117, 22311–22322 (2020).

5. Fischhoff, I. R., Castellanos, A. A., Rodrigues, J. P. G. L. M., Varsani, A. & Han, B. A. Predicting the zoonotic capacity of mammals to transmit SARS-CoV-2. Proc. Biol. Sci. 288, 20211651 (2021).

6. Nerpel, A. et al. SARS-ANI: a global open access dataset of reported SARS-CoV-2 events in animals. Scientific Data 9, 1–13 (2022).

7. Oude Munnink, B. B., et al. Transmission of SARS-CoV-2 on mink farms between humans and mink and back to humans. Science 371, 172–177 (2021).

8. Pickering, B. et al. Divergent SARS-CoV-2 variant emerges in white-tailed deer with deer-to-human transmission. Nature Microbiology 7, 2011–2024 (2022).

9. Yen, H.-L. et al. Transmission of SARS-CoV-2 delta variant (AY.127) from pet hamsters to humans, leading to onward human-to-human transmission: a case study. Lancet 399, 1070 (2022).

10. Feng, A. et al. Transmission of SARS-CoV-2 in free-ranging white-tailed deer in the United States. Nat. Commun. 14, 1–17 (2023).

11. Centers for Disease Prevention and Control. Ecology. https://www.cdc.gov/hantavirus/technical/hps/ecology.html (2019).

12. Machtinger, E. T. & Williams, S. C. Practical Guide to Trapping Peromyscus leucopus (Rodentia: Cricetidae) and Peromyscus maniculatus for Vector and Vector-Borne Pathogen Surveillance and Ecology. J. Insect Sci. 20, 5 (2020).

13. Bosco-Lauth, A. M. et al. Peridomestic Mammal Susceptibility to Severe Acute Respiratory Syndrome Coronavirus 2 Infection. Emerg. Infect. Dis. 27, 2073–2080 (2021).

14. Fagre, A. et al. SARS-CoV-2 infection, neuropathogenesis and transmission among deer mice: Implications for spillback to New World rodents. PLoS Pathog. 17, e1009585 (2021).

15. Griffin, B. D. et al. SARS-CoV-2 infection and transmission in the North American deer mouse. Nat. Commun. 12, 3612 (2021).

16. Lewis, J. et al. SARS-CoV-2 infects multiple species of North American deer mice and causes clinical disease in the California mouse. Front. Virol. 3, 1114827 (2023).

17. Goldberg, A. R., et al. Wildlife exposure to SARS-CoV-2 across a human use gradient. bioRxiv (2023) doi:10.1101/2022.11.04.515237.

18. Shriner, S. A. et al. SARS-CoV-2 Exposure in Escaped Mink, Utah, USA. Emerg. Infect. Dis. 27, 988–990 (2021).

19. Colombo, V. C. et al. SARS-CoV-2 surveillance in Norway rats (Rattus norvegicus) from Antwerp sewer system, Belgium. Transbound. Emerg. Dis. 69, 3016–3021 (2022).

20. Miot, E. F. et al. Surveillance of Rodent Pests for SARS-CoV-2 and Other Coronaviruses, Hong Kong. Emerg. Infect. Dis. 28, 467–470 (2022).

21. Wang, Y. et al. SARS-CoV-2 exposure in Norway rats (Rattus norvegicus) from New York City. mBio 14, (2023).

22. Caserta, L. C. et al. White-tailed deer (Odocoileus virginianus) may serve as a wildlife reservoir for nearly extinct SARS-CoV-2 variants of concern. Proc. Natl. Acad. Sci. U. S. A. 120, e2215067120 (2023).

23. Chandler, J. C. et al. SARS-CoV-2 exposure in wild white-tailed deer (Odocoileus virginianus). Proc. Natl. Acad. Sci. U. S. A. 118, (2021).

24. Hale, V. L. et al. SARS-CoV-2 infection in free-ranging white-tailed deer. Nature 602, 481– 486 (2022).

25. Kotwa, J. D. et al. Genomic and transcriptomic characterization of Delta SARS-CoV-2 infection in free-ranging white-tailed deer (Odocoileus virginianus). bioRxiv (2023) doi:10.1101/2022.01.20.476458.

26. Kuchipudi, S. V. et al. Multiple spillovers from humans and onward transmission of SARS-CoV-2 in white-tailed deer. Proc. Natl. Acad. Sci. U. S. A. 119, (2022).

27. Marques, A. D. et al. Multiple Introductions of SARS-CoV-2 Alpha and Delta Variants into White-Tailed Deer in Pennsylvania. MBio 13, e0210122 (2022).

28. Palermo, P. M., Orbegozo, J., Watts, D. M. & Morrill, J. C. SARS-CoV-2 Neutralizing Antibodies in White-Tailed Deer from Texas. Vector Borne Zoonotic Dis. 22, 62–64 (2022).

29. Roundy, C. M. et al. High Seroprevalence of SARS-CoV-2 in White-Tailed Deer (Odocoileus virginianus) at One of Three Captive Cervid Facilities in Texas. Microbiol Spectr 10, e0057622 (2022).

30. Vandegrift, K. J. et al. SARS-CoV-2 Omicron (B.1.1.529) Infection of Wild White-Tailed Deer in New York City. Viruses 14, (2022).

31. Willgert, K. et al. Transmission history of SARS-CoV-2 in humans and white-tailed deer. Sci. Rep. 12, 12094 (2022).

32. Bevins, S. N. et al. SARS-CoV-2 occurrence in white-tailed deer throughout their range in the conterminous United States. bioRxiv 2023.04.14.533542 (2023) doi:10.1101/2023.04.14.533542.

33. Cool, K. et al. Infection and transmission of ancestral SARS-CoV-2 and its alpha variant in pregnant white-tailed deer. Emerg. Microbes Infect. 11, 95–112 (2022).

34. Martins, M. et al. From Deer-to-Deer: SARS-CoV-2 is efficiently transmitted and presents broad tissue tropism and replication sites in white-tailed deer. PLoS Pathog. 18, e1010197 (2022).

35. Palmer, M. V. et al. Susceptibility of white-tailed deer (Odocoileus virginianus) to SARS-CoV-2. J. Virol. 95, (2021).

36. William, S. C., Linske, M. A. & Stafford, K. C., III. Orally delivered fipronil-laced bait reduces juvenile blacklegged tick (Ixodes scapularis) burdens on wild white-footed mice (Peromyscus leucopus). Ticks Tick Borne Dis. 14, 102189 (2023).

37. Linske, M. A., Williams, S. C. & Li, A. Y. Integrated Tick Management in Guilford, CT: Fipronil-Based Rodent-Targeted Bait Box Deployment Configuration and Peromyscus leucopus (Rodentia: Cricetidae) Abundance Drive Reduction in Tick Burdens. J. Med. Entomol. 59, 591–597 (2021).

38. Williams, S. C., Linske, M. A., DeNicola, A. J., DeNicola, V. L. & Boulanger, J. R. Experimental oral delivery of the systemic acaricide moxidectin to free-ranging white-tailed deer (Artiodactyla: Cervidae) parasitized by Amblyomma americanum (Ixodida: Ixodidae). J. Med. Entomol. 60, 733–741 (2023).

39. Tan, C. W. et al. A SARS-CoV-2 surrogate virus neutralization test based on antibody-mediated blockage of ACE2–spike protein–protein interaction. Nat. Biotechnol. 38, 1073– 1078 (2020).

40. Yale SARS-CoV-2 Genomic Surveillance Initiative. COVIDTracker. https://kphamyale.shinyapps.io/covidtrackerct/.

41. Chitwood, M. H. et al. Reconstructing the course of the COVID-19 epidemic over 2020 for US states and counties: Results of a Bayesian evidence synthesis model. PLoS Comput. Biol. 18, e1010465 (2022).

42. Vogels, C. B. F. et al. Multiplex qPCR discriminates variants of concern to enhance global surveillance of SARS-CoV-2. PLoS Biol. 19, e3001236 (2021).

43. Holbrook, M. G. et al. Updated and Validated Pan-Coronavirus PCR Assay to Detect All Coronavirus Genera. Viruses 13, 599 (2021).

44. Tan, C. C. S. et al. Surveillance of 16 UK native bat species through conservationist networks uncovers coronaviruses with zoonotic potential. Nat. Commun. 14, 3322 (2023).

45. Schoch, C. L. et al. NCBI Taxonomy: a comprehensive update on curation, resources and tools. Database 2020, baaa062 (2020).

46. Lackey, J. A. H. D. G. & Ormiston, B. G. Peromyscus leucopus. Mammalian Species (1985) doi:10.2307/3503904.

47. Wang, W. et al. Antigenic cartography of well-characterized human sera shows SARS-CoV-2 neutralization differences based on infection and vaccination history. Cell Host Microbe 30, 1745–1758.e7 (2022).

48. Collins, C. R. & Kays, R. W. Patterns of Mortality in a Wild Population of White-footed Mice. Northeastern Naturalist 21, 323–336 (2014).

49. Miller, D. H. & Getz, L. L. Comparisons of Population Dynamics of Peromyscus and Clethrionomys in New England. Journal of Mammalogy 58, 1–16 (1977).

50. van Zelm, M. C. Immune memory to SARS-CoV-2 Omicron BA.1 breakthrough infections: To change the vaccine or not? Sci Immunol 7, eabq5901 (2022).

51. Massachusetts Division of Fisheries and Wildlife. Learn about deer. https://www.mass.gov/service-details/learn-about-deer.

52. Hamer, S. A. et al. Persistence of SARS-CoV-2 neutralizing antibodies longer than 13 months in naturally infected, captive white-tailed deer (Odocoileus virginianus), Texas. Emerg. Microbes Infect. 11, 2112–2115 (2022).

53. Iowa State University Extension and Outreach. Mice: Damage Management. https://naturalresources.extension.iastate.edu/encyclopedia/mice-damage-management.

54. Madison, D. M., Hill, J. P. & Gleason, P. E. Seasonality in the Nesting Behavior of Peromyscus leucopus. Am. Midl. Nat. 112, 201 (1984).

55. New Hampshire Fish and Game. White-tailed Deer (Odocoileus virginianus). https://www.wildlife.state.nh.us/wildlife/profiles/deer.html.

56. Mora-Díaz, J. C., Piñeyro, P. E., Houston, E., Zimmerman, J. & Giménez-Lirola, L. G. Porcine Hemagglutinating Encephalomyelitis Virus: A Review. Front Vet Sci 6, 53 (2019).

57. Xu, L. et al. Identification of bovine coronavirus in a Daurian ground squirrel expands the host range of Betacoronavirus 1. Virol. Sin. 38, 321–323 (2023).

58. Hatcher, E. L. et al. Virus Variation Resource - improved response to emergent viral outbreaks. Nucleic Acids Res. 45, D482–D490 (2017).

59. Wang, W. et al. Extensive genetic diversity and host range of rodent-borne coronaviruses. Virus Evol 6, veaa078 (2020).

60. Wang, W. et al. Discovery, diversity and evolution of novel coronaviruses sampled from rodents in China. Virology 474, 19–27 (2015).

61. Embregts, C. W. E. et al. Evaluation of a multi-species SARS-CoV-2 surrogate virus neutralization test. One Health 13, 100313 (2021).

62. Genscript. cPass SARS-CoV-2 Neutralization Antibody Detection Kit Instructions for Use. https://www.genscript.com/gsfiles/techfiles/EUA%2dGenScript%2dcpass%2difu%2epdf?=20220217?1076690102.

63. Berto, A. et al. Detection of potentially novel paramyxovirus and coronavirus viral RNA in bats and rats in the Mekong Delta region of southern Viet Nam. Zoonoses Public Health 65, 30–42 (2018).

64. Ge, X.-Y. et al. Detection of alpha- and betacoronaviruses in rodents from Yunnan, China. Virol. J. 14, 1–11 (2017).

65. Kumakamba, C. et al. Coronavirus surveillance in wildlife from two Congo basin countries detects RNA of multiple species circulating in bats and rodents. PLoS One 16, e0236971 (2021).

66. McIver, D. J. et al. Coronavirus surveillance of wildlife in the Lao People’s Democratic Republic detects viral RNA in rodents. Arch. Virol. 165, 1869–1875 (2020).

67. Monastiri, A. et al. First Coronavirus Active Survey in Rodents From the Canary Islands. Front. Vet. Sci. 8, (2021).

68. Monchatre-Leroy, E. et al. Identification of Alpha and Beta Coronavirus in Wildlife Species in France: Bats, Rodents, Rabbits, and Hedgehogs. Viruses 9, 364 (2017).

69. Tsoleridis, T. et al. Discovery of Novel Alphacoronaviruses in European Rodents and Shrews. Viruses 8, 84 (2016).

70. Alekseev, K. P. et al. Bovine-Like Coronaviruses Isolated from Four Species of Captive Wild Ruminants Are Homologous to Bovine Coronaviruses, Based on Complete Genomic Sequences. J. Virol. 82, 12422 (2008).

71. Sikes, R. S. & the Animal Care and Use Committee of the American Society of Mammalogists. 2016 Guidelines of the American Society of Mammalogists for the use of wild mammals in research and education. J. Mammal. 97, 663–688 (2016).

72. Williams, S. C. & Linske, M. A. Humane Use of Cardiac Puncture for Non-Terminal Phlebotomy of Wild-Caught and Released Peromyscus spp. Animals 10, 826 (2020).

73. Wei, J. et al. Genome-wide CRISPR Screens Reveal Host Factors Critical for SARS-CoV-2 Infection. Cell 184, 76–91.e13 (2021).

74. Fang, Z. et al. Omicron-specific mRNA vaccination alone and as a heterologous booster against SARS-CoV-2. Nat. Commun. 13, 1–12 (2022).

75. Matranga, C. B. et al. Enhanced methods for unbiased deep sequencing of Lassa and Ebola RNA viruses from clinical and biological samples. Genome Biol. 15, 519 (2014).

76. R Core Team. R: A language and environment for statistical computing. (2022).

77. RStudio Team. RStudio: Integrated Development for R. (2020).

78. U.S. Census Bureau. State Population Totals and Components of Change: 2020-2022. (2022).

79. Kahle D, W. H. ggmap: Spatial Visualization with ggplot2. The R Journal 5, 144–161 (2013).

80. Kalantar, K. L. et al. IDseq—An open source cloud-based pipeline and analysis service for metagenomic pathogen detection and monitoring. GigaScience 9, (2020).

81. Saha, S. et al. Unbiased Metagenomic Sequencing for Pediatric Meningitis in Bangladesh Reveals Neuroinvasive Chikungunya Virus Outbreak and Other Unrealized Pathogens. mBio 10, e02877–19 (2019).

82. Ramesh, A. et al. Metagenomic next-generation sequencing of samples from pediatric febrile illness in Tororo, Uganda. PLoS One 14, e0218318 (2019).

83. Langmead, B., Trapnell, C., Pop, M. & Salzberg, S. L. Ultrafast and memory-efficient alignment of short DNA sequences to the human genome. Genome Biol. 10, 1–10 (2009).

84. Katoh, K., Misawa, K., Kuma, K. & Miyata, T. MAFFT: a novel method for rapid multiple sequence alignment based on fast Fourier transform. Nucleic Acids Res. 30, 3059–3066 (2002).

85. Katoh, K. & Standley, D. M. MAFFT Multiple Sequence Alignment Software Version 7: Improvements in Performance and Usability. Mol. Biol. Evol. 30, 772–780 (2013).

86. Guindon, S. et al. New Algorithms and Methods to Estimate Maximum-Likelihood Phylogenies: Assessing the Performance of PhyML 3.0. Syst. Biol. 59, 307–321 (2010).

87. Rambaut, A. FigTree. (2018).

