## Supplementary Information for "Survey of white-footed mice in Connecticut, USA reveals low SARS-CoV-2 seroprevalence and infection with divergent betacoronaviruses"

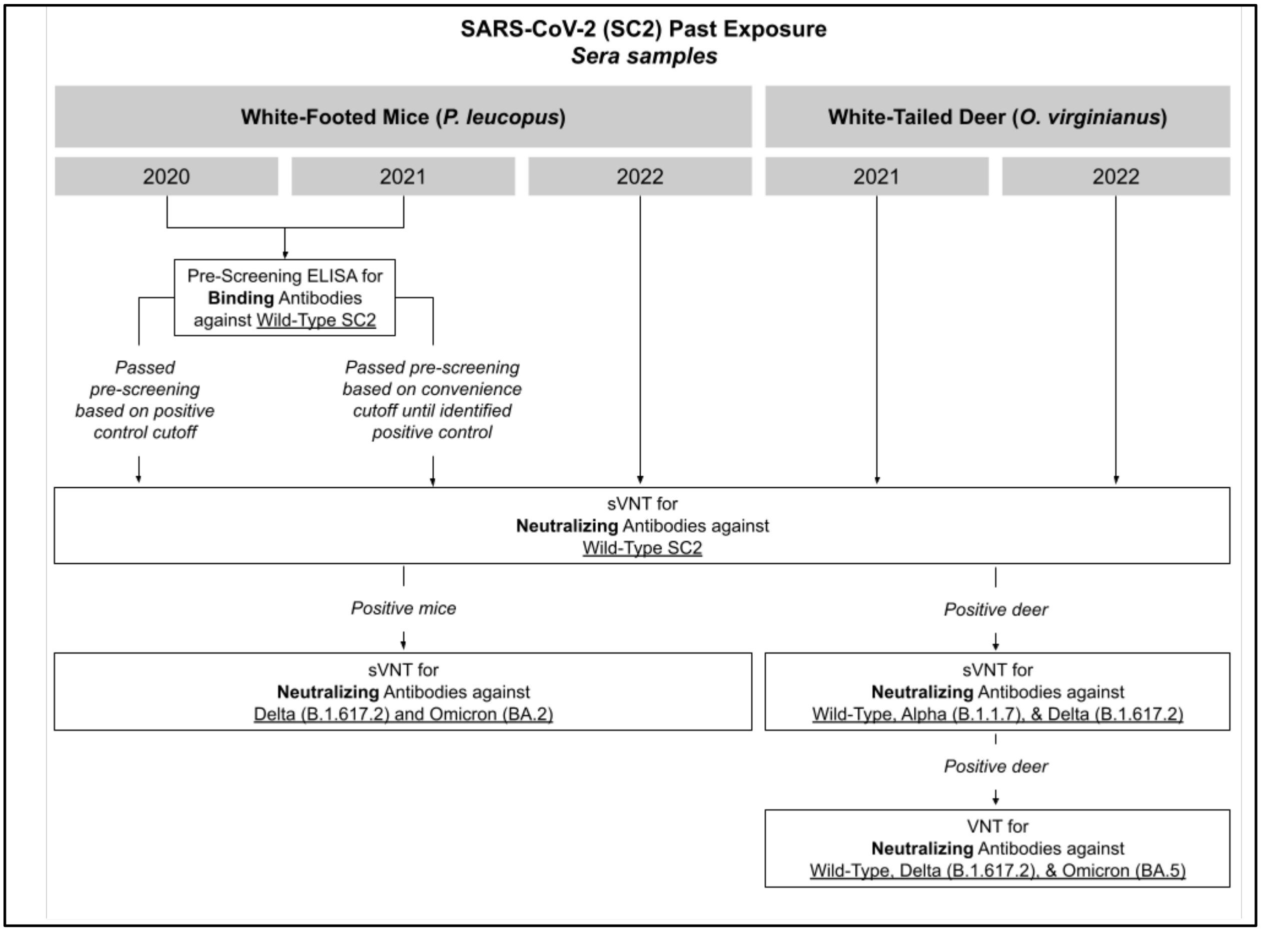


**Supplementary Figure 1: Serological screening flow for white-footed mice and white-tailed deer, related to Figure 1B.** We initially pre-screened 2021 sera samples using an in-house ELISA. Any samples passing a convenience cutoff were screened via the sVNT for wild-type neutralizing antibodies. Once we identified a neutralizing antibody positive white-footed mouse, we included it as a positive control for 2020 sera pre-screening ELISAs and used a positivity cutoff of ≥1x the positive control OD values. We did not pre-screen the 2022 white-footed mice and 2021/2022 deer samples given their relatively higher probability of exposure and smaller number of samples. We tested any wild-type positive samples with sufficient remaining sample volume against various SARS-CoV-2 variants via sVNT.

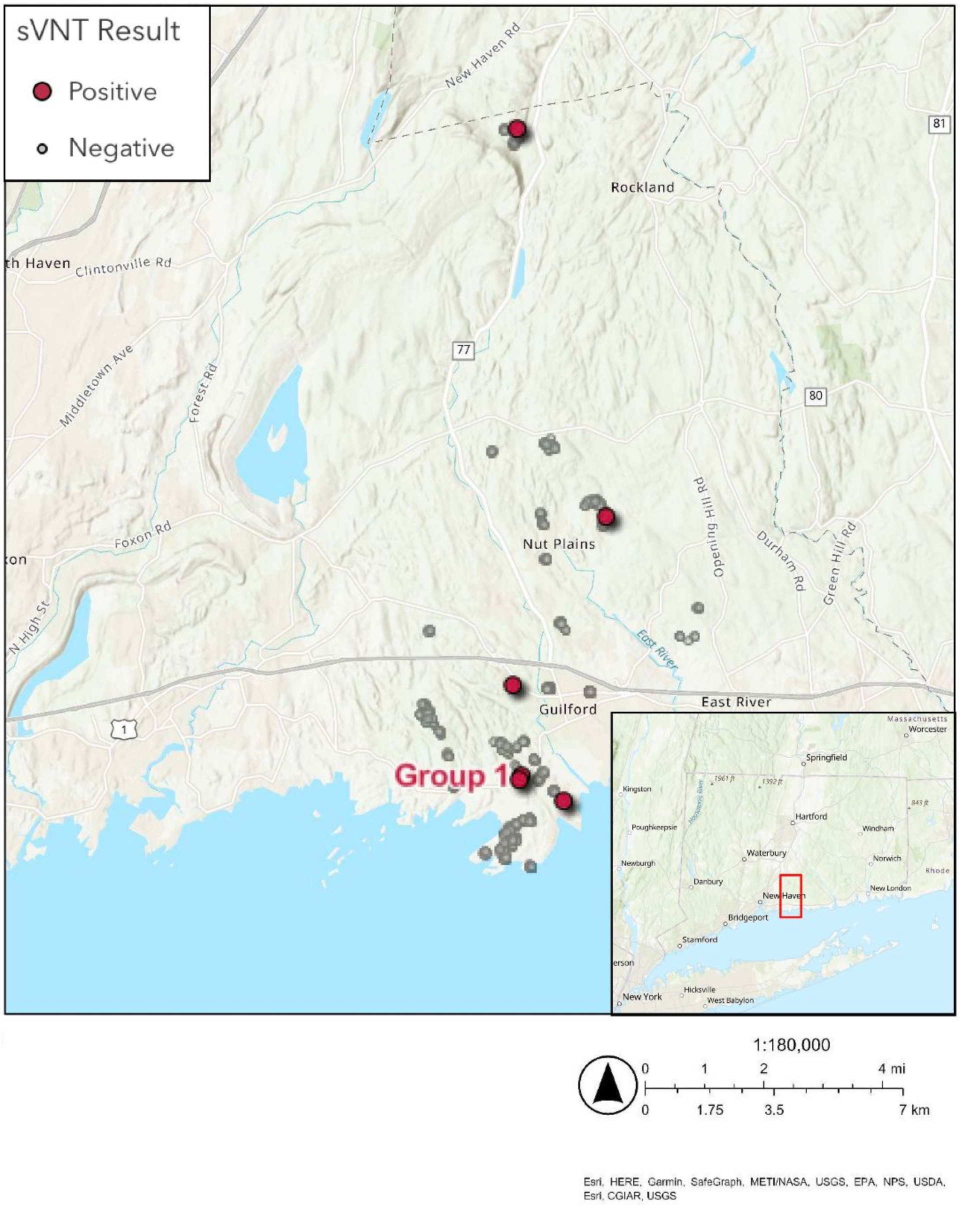


**Supplementary Figure 2: Grouping of wild-type SARS-CoV-2 neutralizing antibody positives among white-footed mice sampled in a residential setting in Guilford, CT, related to Figure 1B.** Map of residential white-footed mice sample collection locations in Guilford, CT by sVNT result. Two neutralizing antibody positive white-footed mice in Group 1 were sampled on the same day from different residential properties separated by approximately 160 meters, placing them within the potential home range of white-footed mice. The remaining positive mice were spatially dispersed. The colors correspond to the wild-type SARS-CoV-2 neutralizing antibody sVNT result for sera samples collected 2020-2022 (see legend). We did not display untested samples and jittered the sample location coordinates for visibility. The inset displays the sampling location within Connecticut.

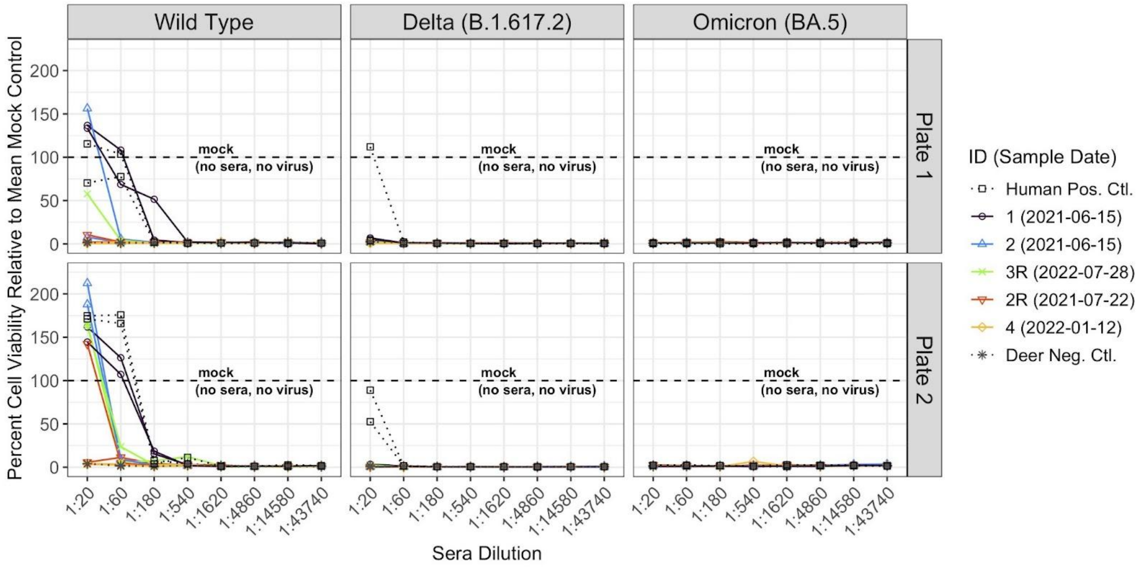


**Supplementary Figure 3: Neutralization of wild-type SARS-CoV-2 by the majority of tested white-tailed deer using a virus neutralization test (VNT), with no cross-neutralization of the Delta (B.1.617.2) or Omicron (BA.5) viral variants, related to Figure 2.** Percent cell viability relative to the mean mock control (no sera, no virus) by SARS-CoV-2 variant using a VNT. We tested previously wild-type SARS-CoV-2 neutralizing antibody positive deer via the sVNT. We tested each sample in duplicate per plate, with two plates prepared by different individuals, against wild-type, Delta (B.1.617.2), and Omicron (BA.5). We included a sVNT negative deer control and a human SARS-CoV-2 positive control, in addition to the mock control. The sample IDs in the legend are in order of descending mean percent cell viability for wild-type SARS-CoV-2. “R” at the end of a sample ID indicates a recaptured animal.

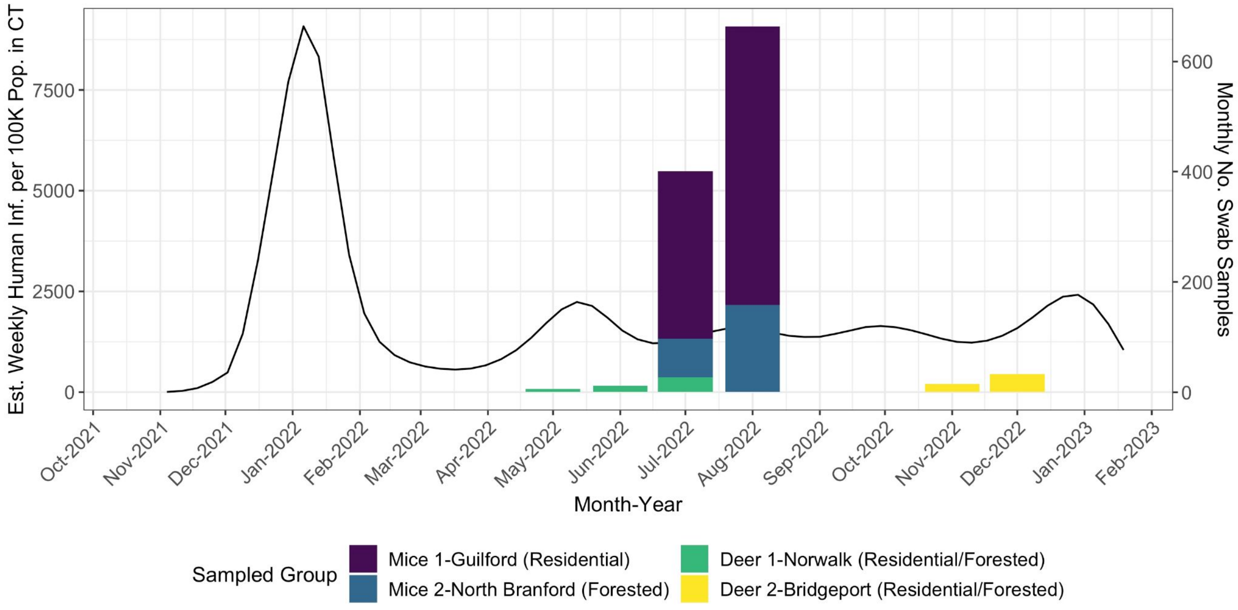


**Supplementary Figure 4: Human SARS-CoV-2 infection context during 2022 swab collection for white-footed mice and white-tailed deer at sampling locations in Connecticut.** Monthly number of total swabs collected for each species and sampling site (bars) and estimated weekly human infections per 100K population in Connecticut (line)[41,78]. We collected two swabs per mouse (oral and anal) and three swabs per deer (oral, anal, and nasal) on each sample collection date. The colors correspond to each species and sampling site.

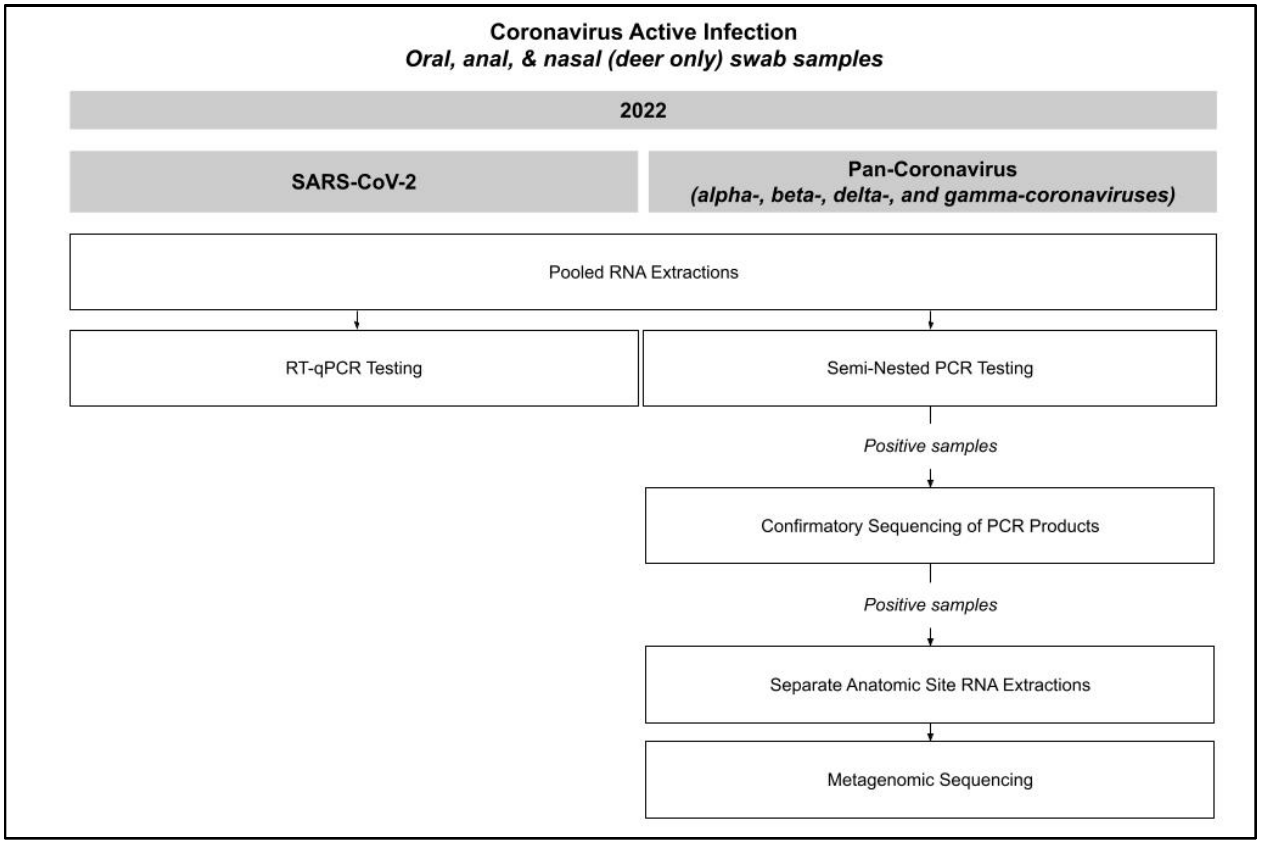


**Supplementary Figure 5: PCR testing and sequencing flow for mice and deer, related to Figure 3 and Supplementary Figure 4.** We pooled swab samples across the anatomic sites per individual and performed RNA extractions. We tested each pooled swab sample via RT-qPCR for SARS-CoV-2 infection and via the semi-nested PCR assay for additional coronaviruses. We did not detect any RT-qPCR SARS-CoV-2 positives. For pan-coronavirus PCR positives, we performed confirmatory sequencing of PCR products. Subsequently, for confirmed positives, we re-extracted the original swabs from each anatomic site (instead of pooling) and performed untargeted metagenomic sequencing.

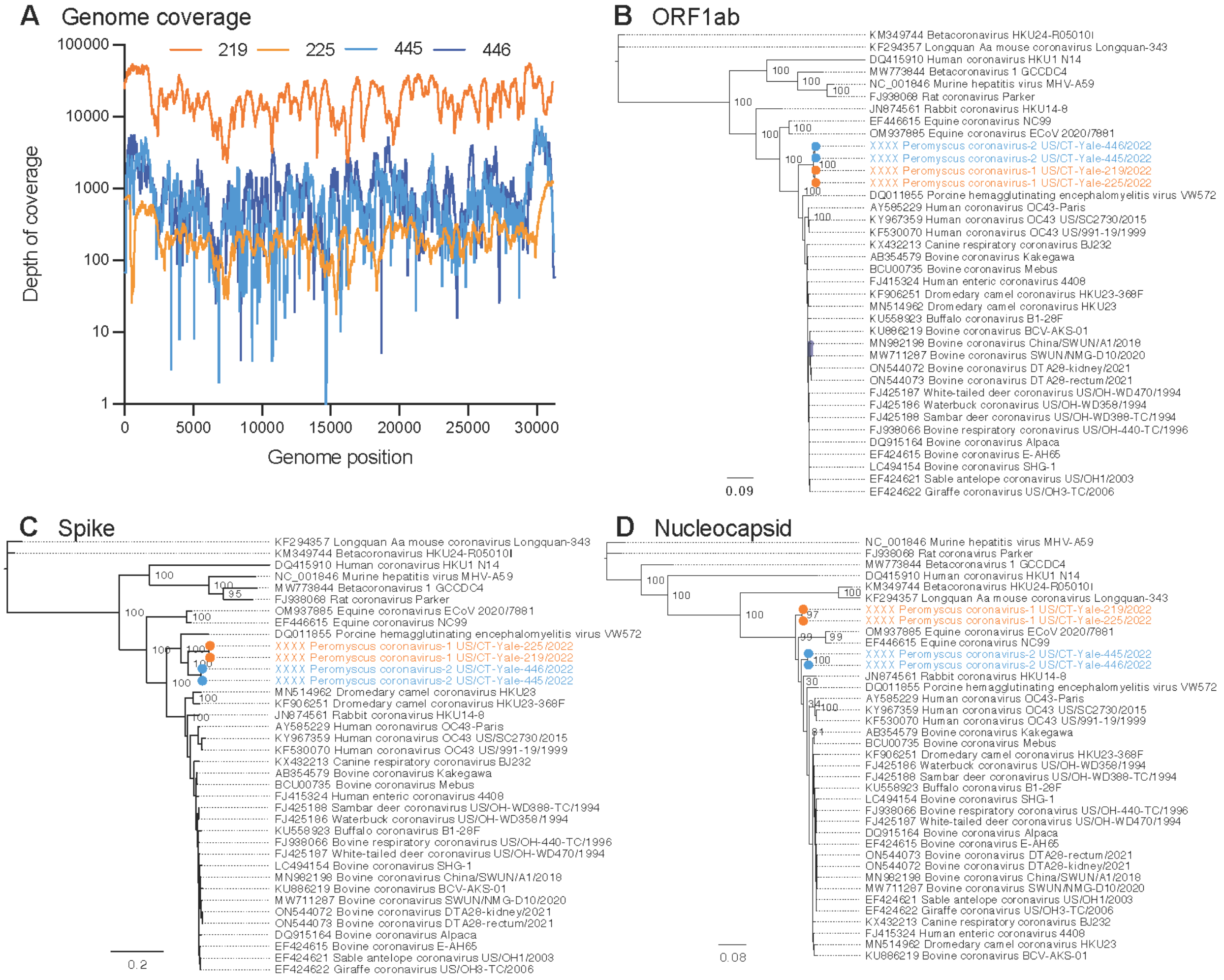


**Supplementary Figure 6: Coverage plot and phylogenetic trees including divergent betacoronaviruses by gene, related to Figure 3B.** A) Depth of sequencing coverage for each of the four sequenced samples containing divergent betacoronaviruses PCoV1 and PCoV2. B-D) Betacoronavirus phylogenetic trees including PCoV1 and PCoV2 genomes by ORF1ab, spike, and nucleocapsid gene, respectively. GenBank accession numbers will be provided when they are available (see Data Availability statement).

| Sequencing ID | Total Reads | CoV Reads | CoV RPM | Percent of Genome | Average Depth |
|---|---|---|---|---|---|
| Peromyscus-coronavirus-1_US/CT-Yale-219/2022 | 622,569,038 | 17,922,056 | 28,787 | 100 | 178,034 |

| | | | | | |
|---|---|---|---|---|---|
| Peromyscus-coronavirus-1_US/CT-Yale-225/2022 | 653,762 | 53,901 | 82,447 | 100 | 255 |
| Peromyscus-coronavirus-2_US/CT-Yale-445/2022 | 46,888,116 | 184,986 | 3,945 | 99.9* | 898 |
| Peromyscus-coronavirus-2_US/CT-Yale-446/2022 | 186,091,038 | 235,777 | 1,267 | 99.6 | 1,117 |

**Gaps filled in with amplicon sequencing.*

**Supplementary Table 1: Metagenomics sequencing information for each identified divergent betacoronavirus genome, related to Figure 3B.** Original animal ear tag number, sequencing ID, total number of reads, total number of betacoronavirus reads, number of reads aligning to the betacoronavirus taxon in the NCBI NR/NT database per million reads sequenced, percent of genome sequenced, and average depth of sequencing.

| GenBank | Strain Name | ORF1 ab | NS | HE | S | NS | NS | NS | E | M | N |
|---|---|---|---|---|---|---|---|---|---|---|---|
| XXXX | Peromyscus-coronavirus-1_US/CT-Yale-219/2022 | 203-21447 | 21457-22292 | 22305-23579 | 23594-27643 | 27633-27997 | 28039-28172 | 28250-28579 | 28566-28820 | 28835-29524 | 29534-30880 |
| XXXX | Peromyscus-coronavirus-1_US/CT-Yale-225/2022 | 203-21456 | 21446-22302 | 22314-23588 | 23603-27652 | 27642-28006 | 28048-28181 | 28259-28588 | 28575-28829 | 28844-29533 | 29543-30889 |
| XXXX | Peromyscus-coronavirus-2_US/CT-Yale-445/2022 | 239-21484 | 21494-22330 | 22342-23616 | 23631-27680 | 27670-28034 | 28076-28209 | 28287-28616 | 28603-28857 | 28872-29558 | 29574-30917 |
| XXXX | Peromyscus-coronavirus-2_US/CT-Yale-446/2022 | 239-21483 | 21493-22329 | 22341-23615 | 23630-27679 | 27699-28033 | 28075-28208 | 28286-28615 | 28602-28856 | 28871-29557 | 29573-30916 |

**Supplementary Table 2: GenBank submission information and gene coding positions for each sequenced divergent betacoronavirus genome, related to Figure 3B.** GenBank accession number, strain name, and gene coding positions. GenBank accession numbers will be provided when they are available (see Data Availability statement).

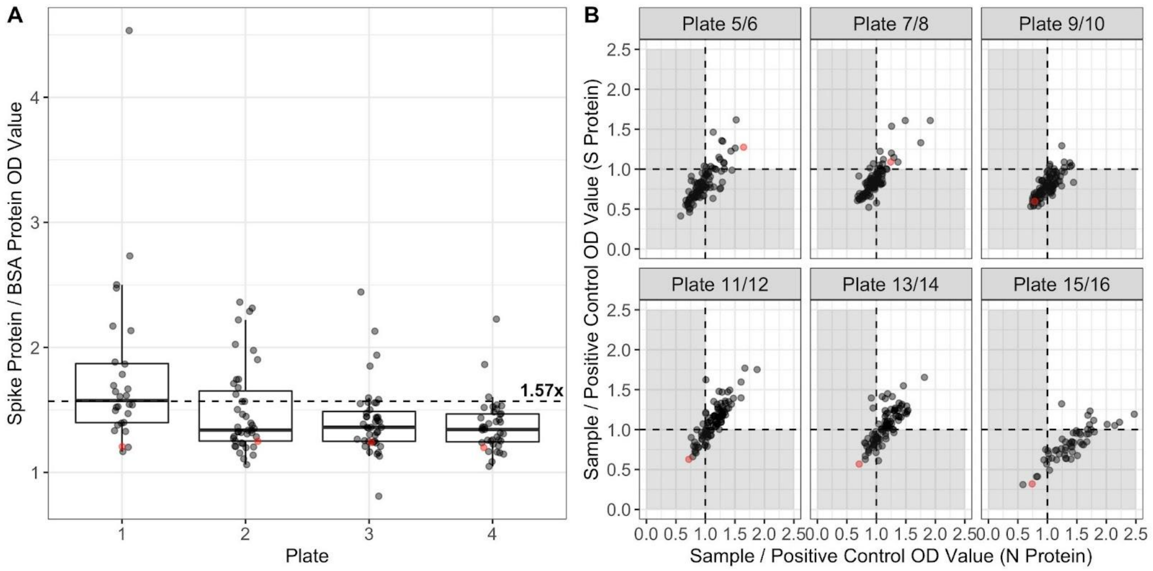


**Supplementary Figure 7: Pre-screening ELISA assay results for wild-type SARS-CoV-2 spike and nucleocapsid protein binding antibodies, related to Figure 1B. A)** 2021 white-footed mouse sera ELISA results (plates 1-4). Initially, lacking a positive white-footed mouse control, we ran each value on a spike (S) protein-coated plate and a BSA protein-coated plate, the latter of which served as a non-SARS-CoV-2 protein comparison. We included any sample with an OD value ≥1.57x for the spike protein compared to BSA protein (a convenience cutoff). **B)** 2020 white-footed mouse sera ELISA results (plates 5-16). After identifying a neutralizing antibody positive white-footed mouse among the 2021 samples, we tested each sample once on a S protein-coated plate and once on a nucleocapsid (N) protein-coated plate. We included any sample with a S and N OD value ≥1x that of the positive control. Red circles indicate the negative control values. We used 2010 white-footed mice sera as negative controls until Plate 9/10, upon which we switched to a sVNT negative 2020 white-footed mice sera control due to elevated values observed with the 2010 white-footed mice negative controls. See Methods for further details. Dashed lines indicate the positivity cutoffs.
